# Systems Analysis of Immune Changes after B-cell Depletion in Autoimmune Multiple Sclerosis

**DOI:** 10.1101/2024.02.07.576204

**Authors:** Jessica Wei, Jeonghyeon Moon, Yoshiaki Yasumizu, Le Zhang, Khadir Radassi, Nicholas Buitrago-Pocasangre, M. Elizabeth Deerhake, Nicolas Strauli, Chun-Wei Chen, Ann Herman, Rosetta Pedotti, Catarina Raposo, Isaiah Yim, Jenna Pappalardo, Erin E. Longbrake, Tomokazu S. Sumida, Pierre-Paul Axisa, David A. Hafler

**Affiliations:** Department of Neurology, Yale School of Medicine, New Haven, CT, USA; Department of Immunobiology, Yale School of Medicine, New Haven, CT, USA; Department of Experimental Immunology, Immunology Frontier Research Center, Osaka University, Suita, Osaka, Japan; Genentech, San Francisco, CA; Roche, Basel, Switzerland; Department of Biomedical Engineering, Yale University, New Haven, CT, USA; Centre de recherches en cancérologie de Toulouse (CRCT), Inserm, Toulouse, France; Broad Institute, Boston, MA, USA

## Abstract

Multiple sclerosis (MS) is a complex genetically mediated autoimmune disease of the central nervous system where anti-CD20-mediated B cell depletion is remarkably effective in the treatment of early disease. While previous studies investigated the effect of B cell depletion on select immune cell subsets using flow cytometry-based methods, the therapeutic impact on patient immune landscape is unknown. In this study, we explored how a therapy-driven “*in vivo perturbation*” modulates the diverse immune landscape by measuring transcriptomic granularity with single-cell RNA sequencing (scRNAseq). We demonstrate that B cell depletion leads to cell type-specific changes in the abundance and function of CSF macrophages and peripheral blood monocytes. Specifically, a CSF-specific macrophage population with an anti-inflammatory transcriptomic signature and peripheral CD16^+^ monocytes increased in frequency post-B cell depletion. This was accompanied by increases in TNFα messenger RNA and protein in monocytes post-B cell depletion, consistent with the finding that anti-TNFα treatment exacerbates autoimmune activity in MS. In parallel, B cell depletion induced changes in peripheral CD4^+^ T cell populations, including increases in the frequency of TIGIT^+^ regulatory T cells and marked decreases in the frequency of myelin peptide loaded-tetramer binding CD4^+^ T cells. Collectively, this study provides an exhaustive transcriptomic map of immunological changes, revealing different mechanisms of action contributing to the high efficacy in B cell depletion treatment of MS.

## Introduction

While B cell depletion is efficacious in the treatment of various autoimmune diseases including rheumatoid arthritis and type 1 diabetes^1,2^, it has conferred remarkable therapeutic benefits in early relapsing remitting multiple sclerosis (MS)^3^. While presumably pathogenic myelin-reactive T cells^4,5^ with loss of regulatory T cell (Treg) function^6–8^ have been established as mechanistic drivers of MS pathogenesis, accumulating evidence directly implicates B cells as key contributors to loss of immune tolerance. The central role of B cells in MS pathophysiology is substantiated by the infiltration of B cells into MS lesions and CSF^9^, the presence of ectopic meningeal B cell follicles adjacent to areas of focal cortical demyelination^10,11^, and the detection of oligoclonal IgG bands in the CSF as a diagnostic marker^12,13^.

Previous studies investigated the effect of B cell depletion on specific immune subsets using hypothesis driven flow cytometry-based methods^14,15^. In this study, we sought to explore the diverse immune landscape and cell states with “*in vivo* perturbation” in patients with MS by measuring immune phenotypes with scRNAseq transcriptomic granularity. B cell depletion in patients was accomplished with systematic removal of naïve and memory B cells through a humanized anti-CD20 antibody, ocrelizumab. We performed 5’ scRNAseq on 18 paired peripheral blood mononuclear cells (PBMC) and paired CSF samples obtained from incident MS patients pre- and post- B cell depletion treatment followed by flow cytometry validation of protein expression. The high dimensional single-cell data set allowed for simultaneous interrogation of the diverse immune populations in patient blood and CSF across tissue types, disease states and treatment status.

Our data revealed increased frequency of a CSF macrophage population that exhibited anti-inflammatory transcriptomic signatures post-B cell depletion therapy. In the periphery, we discovered that CD16^+^ monocytes showed the highest upregulation of transcriptomic TNFα/NF_κ_B signatures after ocrelizumab treatment compared to other cell types. The transcriptional changes were confirmed with increases in TNFα protein in monocytes consistent with the finding that anti- TNFα treatment increases MS disease activity. Moreover, changes in the myeloid compartment were accompanied by increases in TIGIT expressing FoxP3^+^ Tregs and decreases in the frequency of circulating, myelin reactive CD4^+^ T cells. Thus, we defined the immune cell states associated with remission and demonstrate that B cell depletion achieves therapeutic efficacy in early MS through modulating in parallel the innate and adaptive immune compartments.

## Results

### CD14^+^CD68^+^ CSF cells increase in frequency post B cell depletion therapy

All patients were treatment naïve to any immunomodulatory therapy with new onset relapsing remitting MS (Supplementary Table 1). To elucidate the effects of anti-CD20 treatment on the CNS microenvironment, we interrogated five CSF samples from people with relapsing remitting MS (RRMS) pre- and post- B cell depletion treatment and performed an additional comparison with six CSF samples from age-matched healthy donors. All pre-treatment samples were collected from treatment-naïve patients, while post-treatment CSF samples were obtained at different time points: two at 6-month post-treatment, two at 12-month post-treatment, and one at 18-month post-treatment (Supplementary Table 1). After QC of low-quality cells, scRNAseq yielded 60,704 total single cells from 16 CSF samples, including 15,122 cells from six healthy donor samples, 28,493 cells from five MS treatment-naïve samples and 17,089 cells from five MS post- treatment samples (Fig 1a).

**Fig 1:**
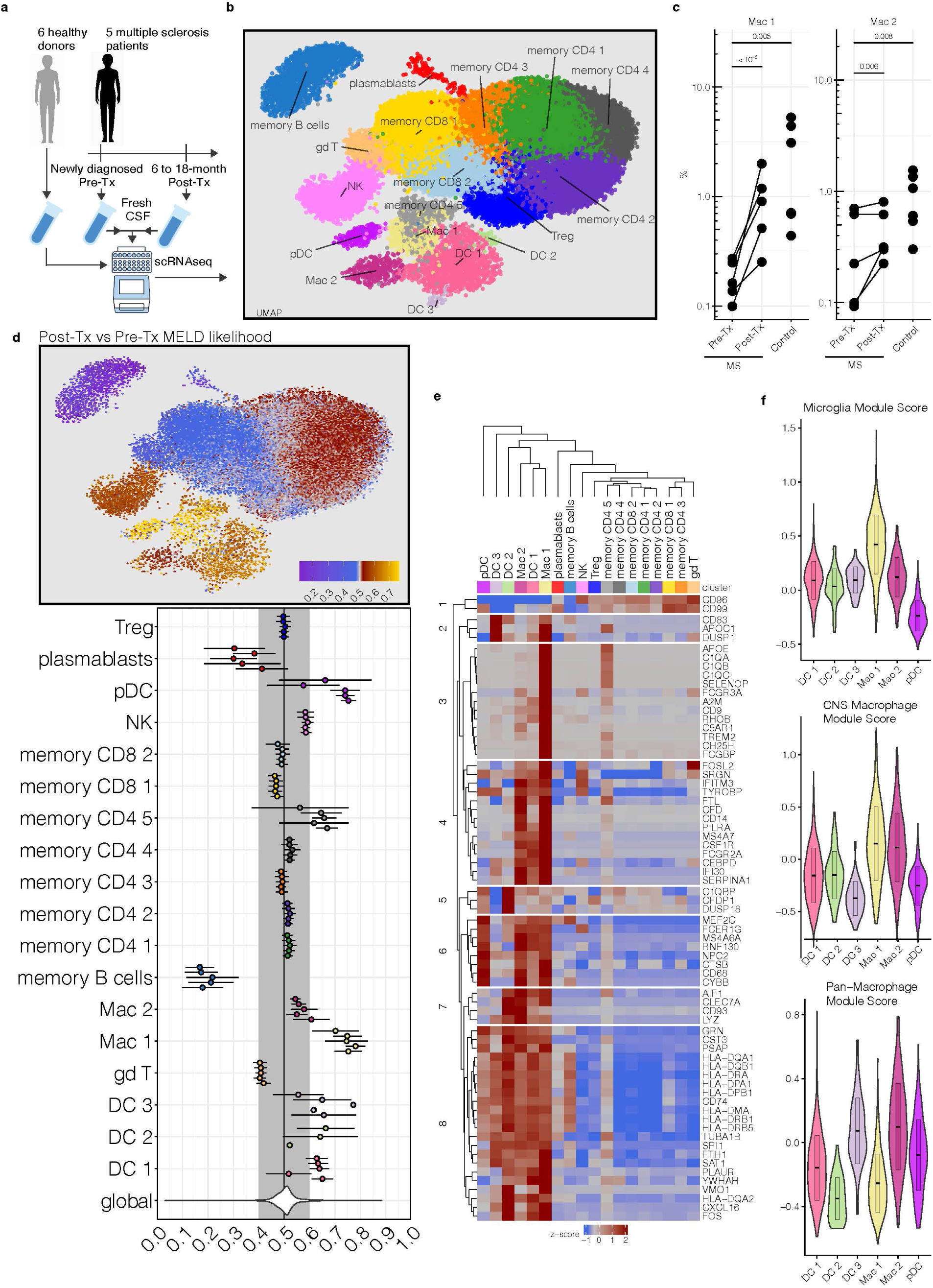
Microglia-like CSF macrophages increase in frequency in MS patients post- B cell depletion therapy. **a,** Healthy donor and MS patient sample collection scheme for scRNAseq analysis (n = 6 healthy donors, n= 5 MS patients pre-treatment, n= 5 matched MS patients post-treatment). **b,** UMAP dimensionality reduction plot of immune cell clusters detected in CSF of healthy donor and MS patients (n= 60,704 single cells, 17 immune cell clusters). **c,** CSF macrophage cluster frequency pre- and post- B cell depletion therapy across all five MS patients. **d,** MELD likelihood enrichment heatmap and patient level summary values (mean+/- SEM, n= 5 MS patients) post- B cell depletion therapy of all immune clusters in the CSF. **e,** Heatmap of myeloid-related genes, showing average expression across all immune cell types. **f,** All myeloid clusters in the CSF scored against microglia, CNS macrophage, and pan- macrophage gene modules.

After normalization and Harmony batch correction (see Methods), CSF cells were classified into 17 clusters (Fig 1b). To assess potential donor and sample variability, the frequencies of all immune populations in the CSF within each patient were profiled. Across all five patients, pre-treatment MS samples had consistently lower cell frequencies in the CD14^+^CD68^+^ myeloid-1 (Mac 1) cluster compared to healthy donors, and the frequencies of this cluster subsequently increased in all patients after B cell depletion treatment (Fig 1c and Supplementary Fig1). In parallel, from all five patients, 1536 B cells were found pre-treatment and only 188 B cells were detected post treatment, as expected due to the mode of action of anti-CD20 therapies. To further identify the immune subset most affected by B cell depletion therapy, we used MELD^16^ to quantify the effect of B cell depletion treatment on all immune cell clusters in the CSF and discovered that the same Mac 1 cluster was the most enriched cell type after treatment (Fig 1d). While the dendritic cell clusters exhibited similar trends of increased frequency post-treatment as the Mac 1 cluster, we prioritized downstream analyses on the Mac 1 subset as it had the highest MELD enrichment score and the least MELD score variability across patients.

### CD14^+^CD68^+^ CSF cells are CSF- specific macrophages with microglia gene signatures

Previous studies reported CD14^+^ CSF cells in various disease settings^17–19^. We aimed to further evaluate the myeloid transcriptomic signatures in our patient CSF samples to define the CD14^+^CD68^+^ myeloid clusters. Mac 1 cluster exhibited high levels of microglia (*TREM2, SPI1, CD68, MEF2C*) and monocyte (*CD14, FCGR3A, CSF1R, HLA-DRA*) transcriptomic signatures, while lacking hallmark microglial genes such as *SALL1*, *P2RY12, FCRLS* and *TMEM119*. In addition to pan-macrophage lineage markers *HLA-DR* and *CD14*, Mac 1 cluster also exhibited high expression of *APOE, CSF1R*, and genes that are expressed in extra parenchymal CSF macrophages such as *CST, TGFB1, MS4A7, LYZ, CLEC7A*. The mixture of microglial-like and monocytic genes and the abundant expression of *C1Q* and *HLA* class II genes classified this cluster as CSF-specific macrophages, distinct from monocytes, microglia and macrophages in other CNS tissues (Fig 1e)^20^. Interestingly, memory CD4 cluster 5 exhibited a similar (albeit muted) signature, leading to the initial co-clustering of those two clusters despite coming from two separate lineages (Supplementary Fig 2a, b and Methods).

To further delineate the CSF macrophage subset, we computed pan-macrophage (*CD44, CCR2, CD45, CD206, CD163, CD274, CD169, MYB*), CNS macrophage (*TGFBI, MS4A7, MS4A6C, LYZ2, CD163, P2RX7, CST, CLEC7A*), and microglia (*P2Y12R, TMEM119, TREM2, CD115, CD172A, CD91, SPI1, FCRLS, SALL1, HEXB, SIGLECH, SLC2A5*) subset-specific gene modules to evaluate module scores on all myeloid clusters in the CSF. The post-treatment enriched Mac 1 cluster scored the highest on the microglia module compared to other myeloid clusters. In contrast, CD14^+^CD68^+^ myeloid-2 (Mac 2) cluster scored higher on the pan-macrophage module compared to others (Fig 1f). Collectively, these transcriptomic signatures reflect the phenotypic diversity of the macrophage compartment in the CSF, and that anti-CD20 treatment increased the frequency of a specific subset of microglia-like macrophages within the CSF.

### B cell depletion induces an anti-inflammatory phenotype in CSF macrophages

With the recent identification of CSF-specific macrophages^20^ and the limited availability of patient CSF samples, the role of CSF macrophages in homeostasis and during MS pathogenesis remains unclear. To better understand the treatment effect of B cell depletion on CSF macrophage immunophenotype, we performed differential gene expression (DE) analysis of pre- and post- treatment MS samples along with healthy donor CSF (Fig 2a). In the enriched Mac 1 cluster, hierarchical clustering on differentially expressed genes in MS pre-treatment vs healthy donor cells revealed extensive changes, segregating cells from the two groups and highlighting alterations in MS patients (Supplementary Fig 3a). In contrast, only 82 genes were differentially expressed between MS pre- and post- treatment CSF macrophages, therefore grouping CSF macrophages from patients together by hierarchical clustering despite treatment status (Supplementary Fig 3b). Among the 60 genes upregulated after B cell depletion are *EGR1* that represses macrophage activation^21^, *ZFP36* and *ZFP36L1* that modulate post-transcriptional regulation of immune responses and oxidative phosphorylation genes (*NDUFA13, UQCRB, NDUFB1, ATP5F1E*) that are associated with anti-inflammatory macrophages^22,23^. In addition, genes involved in migration and endocytosis (*SLC11A1, ITGB2, CSF1R, DAB2, NUMB, RAB11A, ANXA1, APOE*) were downregulated after B cell depletion (Fig 2a).

**Fig 2:**
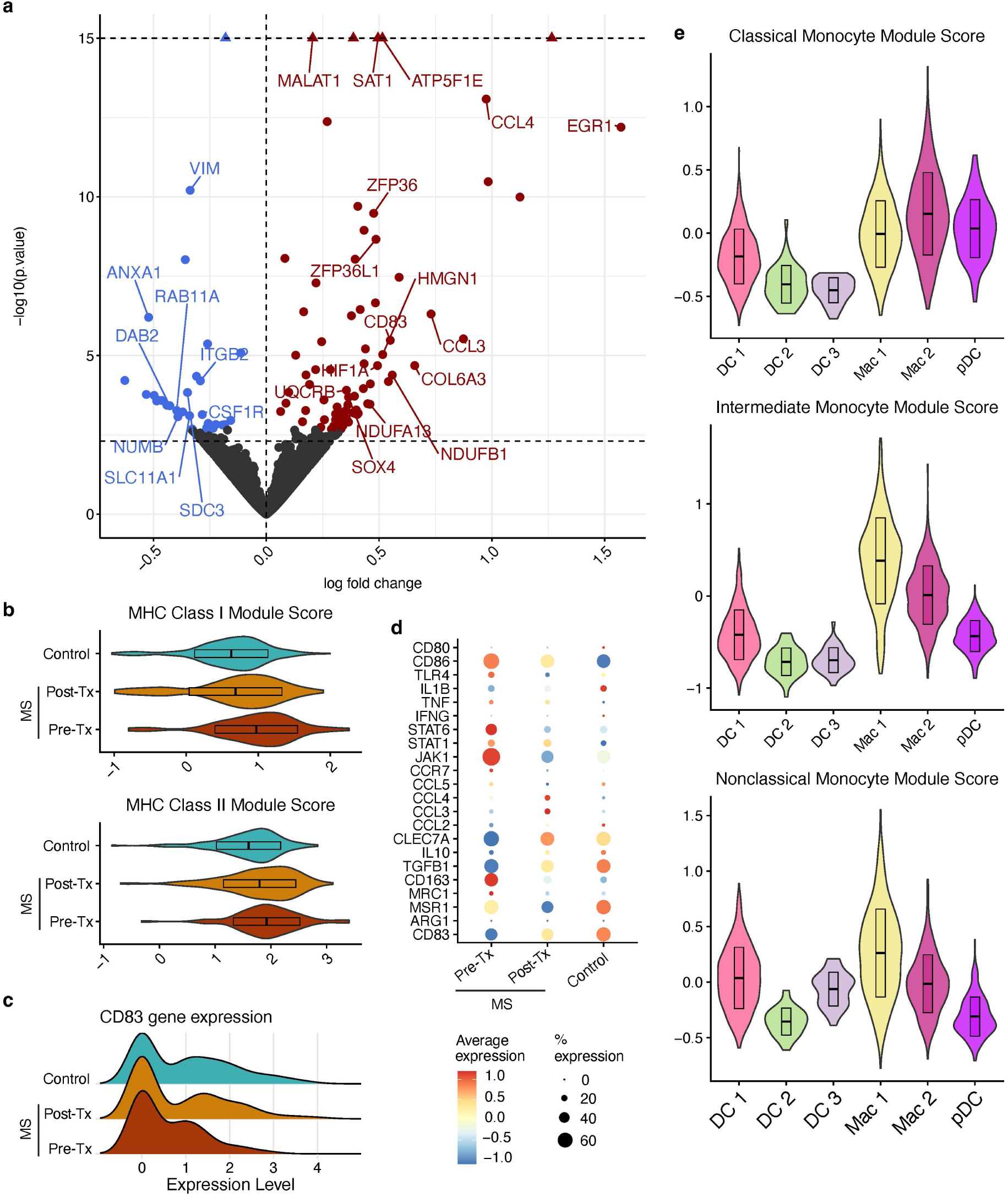
Enriched CSF macrophages present anti-inflammatory phenotype in MS patients post- B cell depletion therapy. Gene expression analyses of CSF macrophages were performed by comparing cells from healthy donors (n=6) and pre- and post- B cell depletion therapy MS patient samples (n=5). **a,** Volcano plot depicting differentially expressed genes in the Mac 1 cluster from MS patients pre- and post- B cell depletion. Blue: downregulated post-treatment, red: upregulated post-treatment. -log10(p value) > 15 are capped to facilitate visualization (depicted as triangles). **b,** MHC class I and class II gene module scores of healthy donor, MS pre-treatment, and MS post-treatment Mac 1 cells. **c,** CD83 expression of healthy donor and MS patient Mac 1 cells. **d,** Dot plot depicting myeloid inflammatory and anti-inflammatory gene expression in healthy donors and MS patients pre- and post- treatment. **e,** Gene module scores of all CSF myeloid clusters against peripheral monocyte gene signatures. Top: classical monocyte module score, middle: intermediate monocyte module score, bottom: nonclassical monocyte module score.

We observed increases in HLA class I and class II mRNA expression in Mac 1 cells from baseline MS patient CSF as compared to healthy CSF Mac 1 cells. These increases in MHC expression from patients with MS decreased after B cell depletion (Fig 2b). In contrast, CD83, a macrophage immune checkpoint marker that contributes to the resolution of inflammation and can induce Tregs in experimental models of MS^24–26^, was downregulated in MS pre-treatment cells, and recovered to healthy donor levels post- B cell depletion treatment (Fig 2c). We next applied the classical and alternative macrophage activation paradigm to delineate the neuroinflammatory state of CSF macrophages. Post- treatment Mac 1 cells exhibited transcriptomic downregulation in pro-inflammatory programming (*CCR7, JAK1, STAT1, IL1B, TNF, TLR4, CD86*) and upregulation of anti-inflammatory genes (*IL-10, TGFB, CLEC7A*) In addition, we observed decrease of macrophage scavenger receptors (*CD163, MRC1, MSR1*) (Fig 2d). Taken together, the decreases in MHC class I and class II gene module scores suggest that B cell depletion reduces the probability of T cell activation through CSF macrophage antigen presentation. The increased expression in *CD83, TGFB, IL10*, and the oxidative phosphorylation pathway in CSF macrophages post-treatment suggests that B cell depletion therapy promotes the resolution of inflammatory phenotype in CSF macrophages to restore homeostasis. Lastly, we computed transcriptomic signature scores based on peripheral monocytic gene modules and applied them to the CSF myeloid populations. We found that the enriched Mac 1 cluster scored the highest on the intermediate (*HLADR, CD14, CD11C, CD68, FCGR3A, CX3CR1, CSF1R, TLR4*) and nonclassical monocyte (*FCGR3A, CX3CR1, SLAN, CSF1R, CXCR1, CXCR4*) gene modules, whereas Mac 2 scored higher on the classical monocyte (*CD14, CCR2, CCR5, SELL, CD36, CD33, CD64*) gene module (Fig 2e). Thus, the transcriptomic resemblance of CSF macrophages to intermediate monocytes prompted us to investigate whether B cell depletion therapy modulates intermediate monocyte frequency in the periphery.

### Increased abundance of CD16^+^ monocytes post B cell depletion in MS patient PBMC

We next investigated whether alterations observed in the CSF post-B cell depletion are recapitulated in peripheral blood. We performed immune profiling with cryopreserved PBMC using scRNAseq from 18 newly diagnosed treatment-naïve patients with relapsing remitting RRMS whose samples were collected for both pre- and 6 months post-treatment (Supplementary Table 1, Fig 3a). In our unsupervised analysis we retained 38 communities, which we assigned to coarse-grained immune cell types of interest (naïve T cells, memory CD4^+^ T cells, cytotoxic lymphocytes, B cells, myeloid cells), and 18 fine-grained cell-types, as described in Supplementary Fig 4a, b and Fig 1b. We classified communities into main lineages based on marker gene inspection and scoring cells against reference datasets using singleR software (Supplementary Fig 4c)^27^.

**Fig 3:**
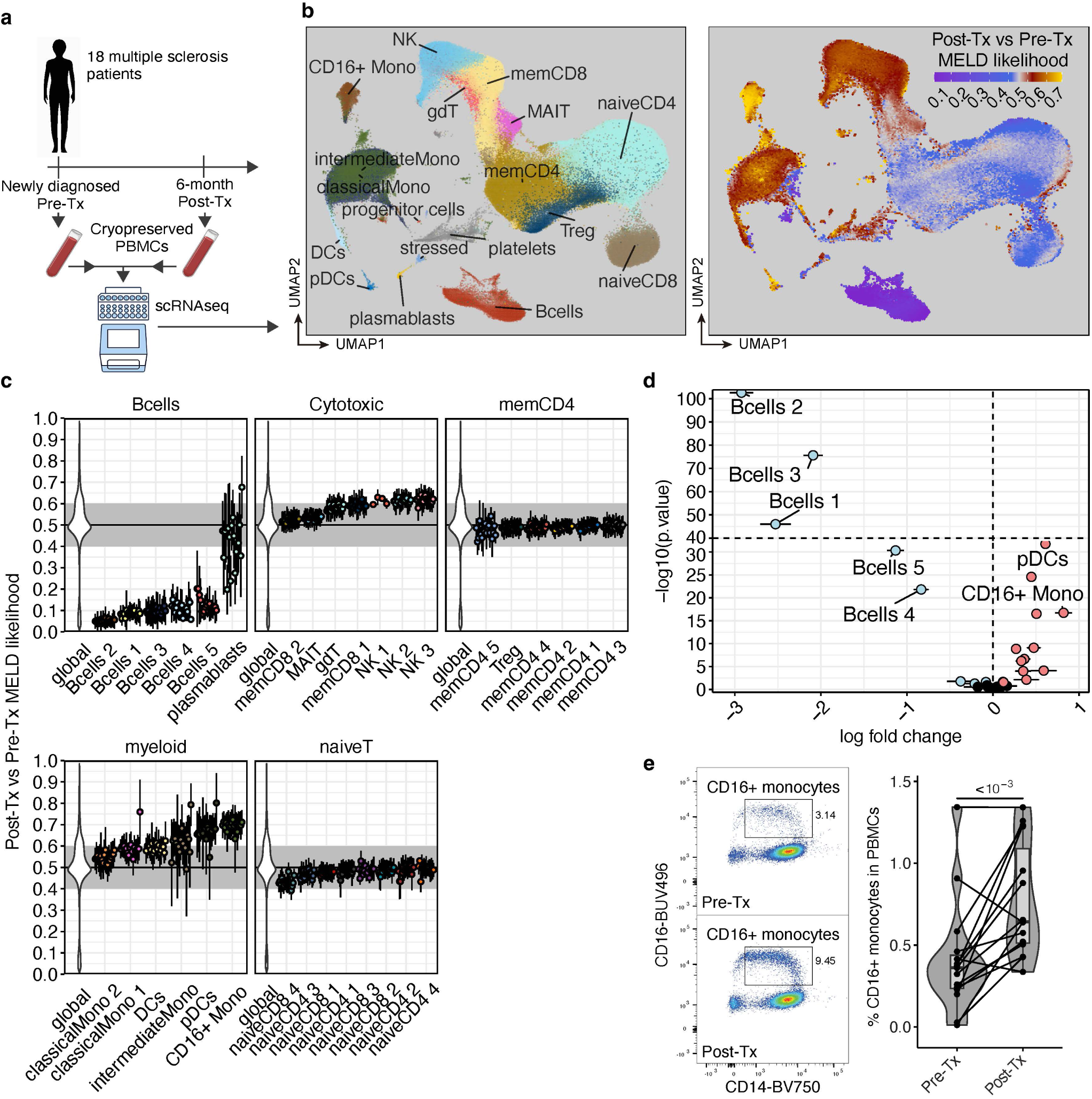
Increased CD16^+^ monocytes abundance after anti-CD20 treatment. a, experimental design of pre- and post-treatment (anti-CD20) PBMC samples from MS patients (n=18) for droplet-based scRNAseq using 10x genomics platform. **b,** UMAP of annotated cell types (left) and overlayed MELD likelihoods for post-treatment status (right). **c,** MELD likelihood patient level summary values (mean+/- SEM) per fine-grained clusters and main cell types. **d,** fine-grained community frequency changes post-treatment (log fold change mean estimate +/-SE from beta regression, see methods). **E,** Flow cytometry validation of CD16^+^ monocyte frequency changes in MS patients PBMCs.

Using MELD, we observed computed differences in abundance between the pre- and post-treatment samples. As expected, ocrelizumab treatment significantly decreased the number of B cells except for plasmablasts, which are known to downregulate CD20 that leads to loss of sensitivity to ocrelizumab-mediated depletion (Fig 3b). Given our CSF data, we next focused our attention on myeloid cells. We observed cellular enrichment in two myeloid clusters: CD16^+^ monocytes and plasmacytoid dendritic cells (pDCs) (Fig 3c). We confirmed these changes by formally testing for variations in frequency across all donors and observed that the increased frequency of CD16^+^ monocytes was conserved across donors (Fig 3d and Supplementary Fig 5a). The CD16^+^ monocytes cluster was the only cluster with detectable levels of *FCGR3A*, the gene encoding CD16 (Supplementary Fig 5b and d), and displayed markers associated with intermediate and non-classical monocytes. Scoring against the Monaco Immune reference with singleR revealed strong enrichment in intermediate and non-classical reference transcriptomes (Supplementary Fig 5c). To confirm this change in abundance we measured circulating frequencies of various monocyte subpopulations using flow cytometry (Supplementary Fig 6a). While there was no significant difference after B cell depletion in CD14^+^CD16^-^ monocytes, there was a significant increase in CD14^+^CD16^+^ monocytes with treatment (Fig 3e).

### Increased activation and TNFα production in CD16^+^ monocytes post B cell depletion

We next determined whether CD16^+^ monocytes harbor an altered transcriptomic state after B cell depletion. We computed differentially expressed genes while controlling for inter-individual variation using a generalized linear mixed model (glmm, as implemented in NEBULA^28^.

We observed an increased expression of soluble molecules (*CCL5, CXCL8, TNFA*), surface receptors (*CD83, ITGB2, ITGA2B*), and NF_κ_B signaling pathway (*TRAF1, NFKB2, REL, RELB, NFKBIA*) (Fig 4a). CD16^+^ monocytes showed downregulation in major transcription factors (*RXRA, IKZF1, KLF4, IRF4*) and CD81, a marker that facilitates monocytes homing to the CNS in EAE^29^. Given the activated transcriptomic signature observed in CD16^+^ monocytes, we validated changes at the protein level and observed downregulation of CD81, and upregulation of monocyte activation marker HLA-DR using flow cytometry (Fig 4b).

**Fig 4:**
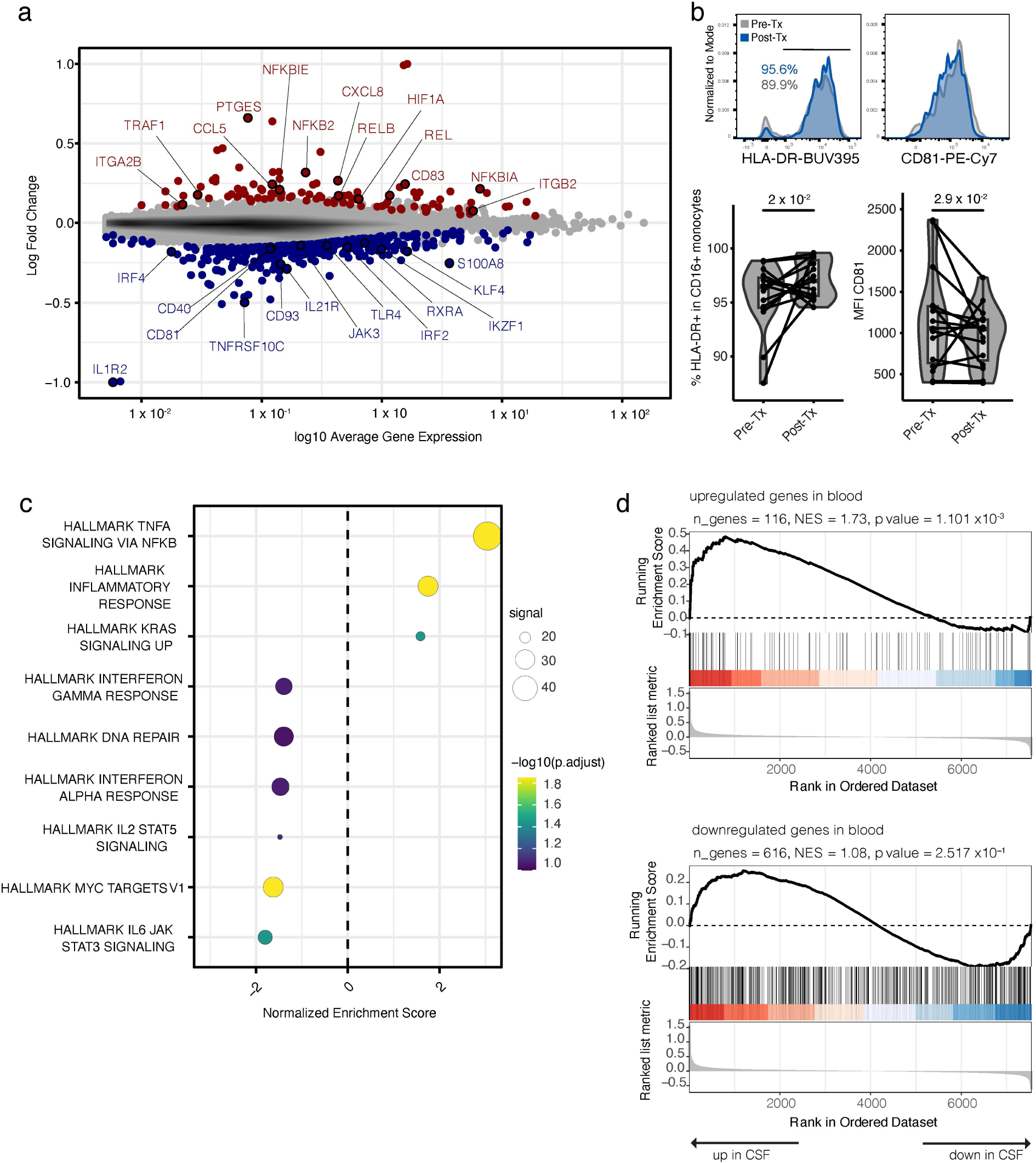
Differential gene expression in CD16^+^ monocyte post-treatment. A, Mean abundance (MA) plot of gene expression changes, differentially expressed genes (DGE) are highlighted in red (upregulated post-treatment), or blue (downregulated post-treatment). **B,** Flow cytometry of HLA-DR and CD81 expression in CD16^+^ monocytes (n=16). **C,** GeneSet Enrichment Analysis (GSEA) using the Hallmark genesets in CD16^+^ monocytes. **D,** Custom GSEA analysis of PBMC monocytes post-treatment signature genesets (up- and downregulated genes) tested on CSF macrophage Mac 1 dataset (From Fig 1).

Given the presence of differentially expressed genes from the NF_κ_B signaling pathway, we next performed GSEA analysis in the enriched CD16^+^ monocyte cluster using the Hallmark gene sets collection. Consistent with these findings, we observed enrichment in TNF-NF_κ_B pathway (Fig 4c), as well as downregulation related to JAK-STAT signaling gene sets (IL-2, STAT5, IL-6, JAK, STAT3, interferon alpha and interferon gamma signaling) post-treatment. Finally, to determine whether these pathway modulations can also be observed in CSF macrophages (described in Figure 2), we created custom gene sets based on PBMC DE and ran GSEA on the enriched Mac 1 CSF macrophages to test for gene signature enrichment. We observed a positive enrichment of blood-upregulated genes in the CSF, while blood-downregulated genes showed no significant enrichment in the CNS (Fig 4d). These data suggest that while B cell depletion therapy upregulates similar biological signatures in the CNS and periphery, the treatment-mediated *in vivo* perturbation of biological pathways is tissue-dependent and informed by the local environment.

### B cell depletion induces ubiquitous response to TNFα in PBMC

We next assessed whether the observed upregulation of TNF-NF_κ_B pathway post- treatment is cell-type specific. We computed differentially expressed genes for pre- vs post- B cell depletion in all clusters and used DE results to run GSEA analysis. We observed TNF-NF_κ_B pathway activation across most immune cell types post-ocrelizumab treatment (Fig 5a). However, the downregulation of JAK-STAT related pathways was restricted to CD16^+^ monocytes, and the remaining or repopulating B cells post-treatment showed increased expression of various gene sets related to cell survival (P53 pathway, apoptosis), metabolism (cholesterol homeostasis) and JAK-STAT signaling (IL2 STAT5 signaling, IL6 JAK STAT3 signaling). Interestingly, clustering communities based on GSEA leading edge genes similarity showed that the NF_κ_B signaling enrichment was most similar between B cells and myeloid cells while T lymphocytes formed a separate cluster, suggesting that transcriptomic responses to NF_κ_B signaling differ between those lineages (Fig 5b).

**Fig 5:**
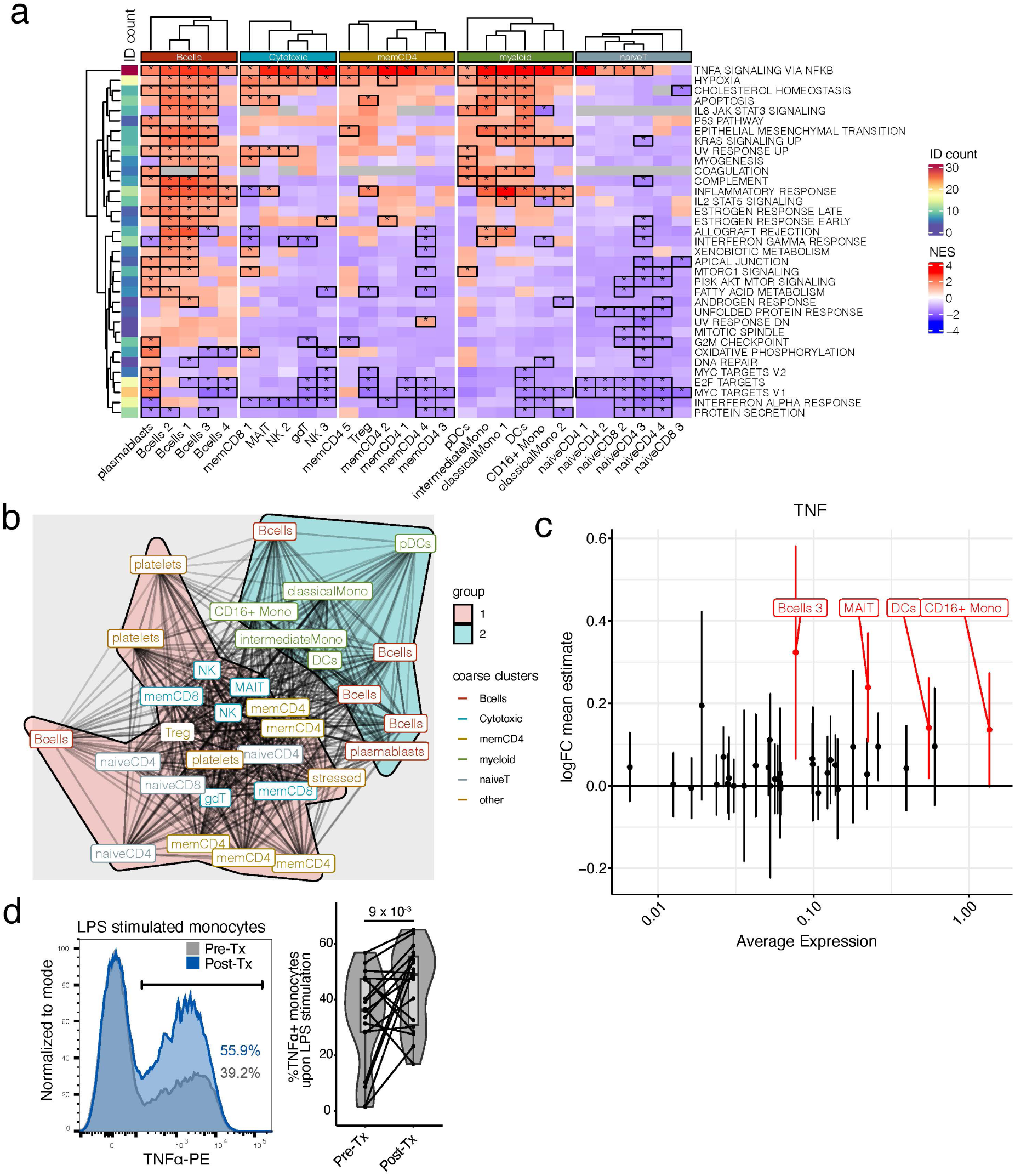
Geneset enrichment analysis (GSEA) of anti-CD20 gene expression alterations across cell types. **a,** Heatmap of normalized enrichment scores (NES) from post-treatment GSEA analyses run for each cluster shows ubiquitous increase in TNFα-NF_κ_B pathway. Differentially enriched genesets are highlighted with a *. “ID count” depicts the number of times a geneset is found enriched across communities. **b,** Overlap graph-analysis of leading edge genes for the “TNFα signaling via NF_κ_B” geneset across cell types highlights two sets of signatures: B and myeloid cells vs. T cells. **c,** Pre- and post- treatment fold change of *TNFA* transcript across clusters (differential expression is highlighted in red). **d,** *In vitro* validation of TNFα upregulation pre- and post- B cell depletion at the protein level in MS patient monocytes (n= 18) by intracellular flow cytometry staining after LPS stimulation.

Despite the ubiquitous TNF-NF_κ_B pathway activation across cell types, CD16^+^ monocytes showed the highest pre-treatment transcriptomic expression of TNFα and sustained the upregulated expression after treatment (Fig 5c). We also detected an increased expression of TNFα in B cells cluster 3, MAIT, and DC clusters. To confirm changes in TNFα expression at the protein level, we enriched CD14^+^ or CD16^+^ monocytes from cryopreserved PBMCs and showed that LPS-stimulated monocytes from MS patients expressed more TNFα post B cell depletion (Fig 5d). Increases of TNFα production by LPS-stimulated CD14^+^ cells was similarly observed in MS patients treated with ocrelizumab by another group^30^. However, there was no significant difference in TNFα ELISA using culture supernatant (data not shown).

### B cell depletion therapy reprograms the CD4^+^ T helper cell compartment in both CSF and PBMC

Though CD4^+^ T cells are thought to play a central role in MS pathophysiology, limited changes were observed in the CD4^+^ T cell compartment using singleR, MonacoImmune, and manual cellular annotation methods. Therefore, we applied a recently developed framework that captures a more granular classification and qualitative assessment of CD4^+^ T cells based on scRNAseq data (Fig 6a)^31^. This novel framework assigned CD4 T cells into five major clusters (cluster Layer 1 or L1) and 18 minor clusters (cluster L2) by Symphony reference mapping (Fig 6b, Supplementary Fig 7a and 8a), and measured the activities of 12 pre-defined transcriptomic gene programs of CD4^+^ T cells using non-negative matrix factorization projection (NMFproj).

**Fig 6:**
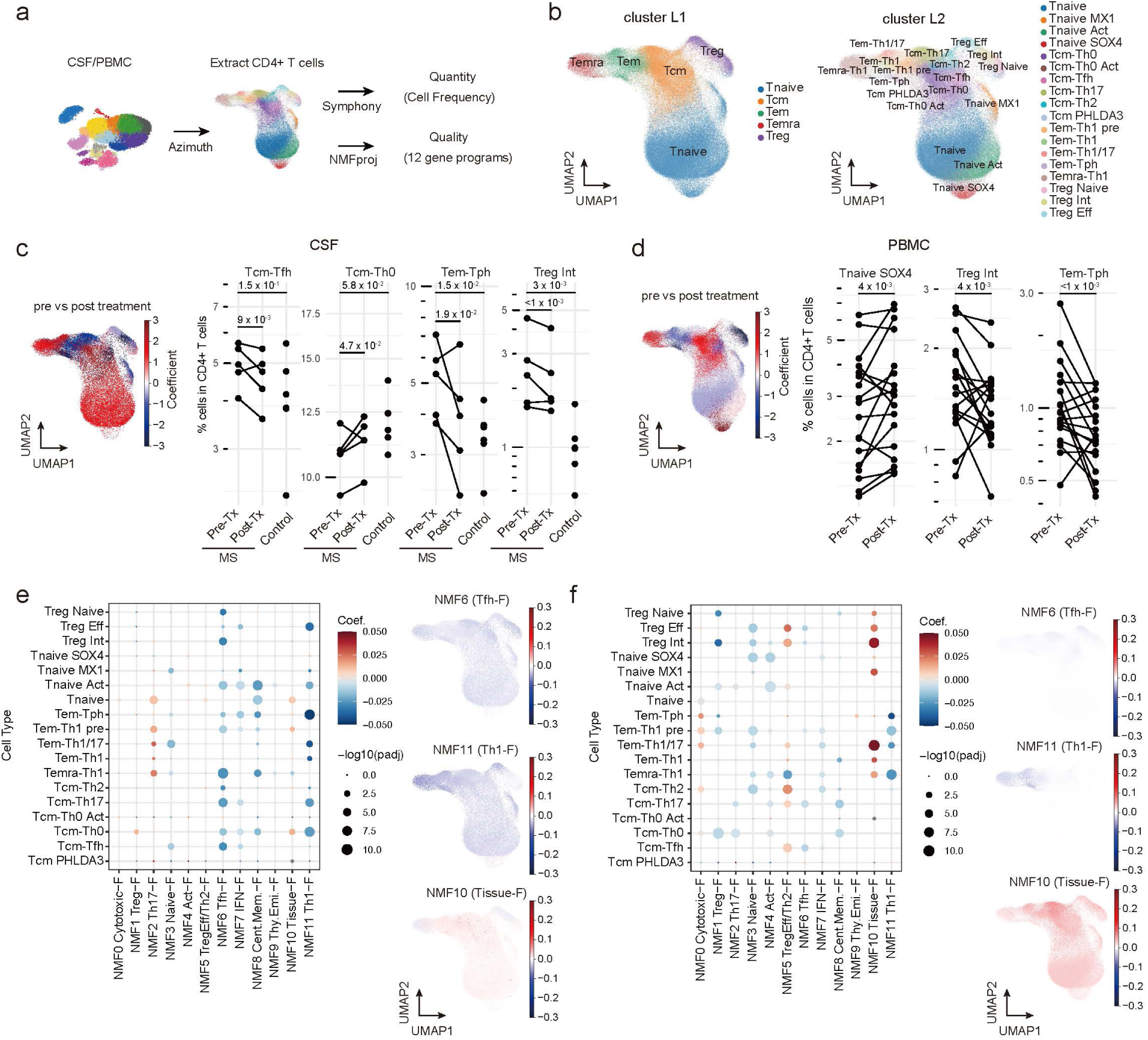
Detailed Analysis of CD4^+^ T Cell Alterations Following Anti-CD20 Treatment. **a,** Schematic illustration of the analysis of CD4^+^ T cells using a reference mapping and NMFproj. From CSF and PBMC samples, CD4^+^ T cells were extracted using Azimuth, and detailed CD4^+^ T clusters were predicted using Symphony. The 12 gene programs were calculated using NMFproj. **b,** Inferred CD4^+^ T cell clusters on UMAP plot. The clusters were assigned to either a major cluster (L1) or a detailed cluster (L2) level. **c, d,** Cell frequency changes after anti-CD20 treatment in CSF **(c)** and PBMC **(d)**. Coefficients of cell frequency change per cluster L2 quantified GLM (method) are visualized on the UMAP plot (left). The populations with cell frequency increases post-B cell depletion treatment are shown in red. CD4^+^ T cluster frequency pre- and post-B cell depletion therapy (right). Significantly altered clusters are shown. See Supplementary Figure 8c and 9c for additional details. **e, f,** Alterations of gene programs extracted by NMFproj after anti- CD20 treatment in CSF **(e)** and PBMC **(f)**. Dot plots depicting NMF cell feature changes in each cell type (left). Dot colors show coefficients, and sizes show the significance of GLM (method). The coefficient of gene program change per cluster for some gene programs was shown on UMAP plots (right). Annotations and representative genes of gene programs are following; NMF0 (Cytotoxic-Feature or Cytotoxic-F; *GZMB*, *CX3CR1*), NMF1 (Treg-F; *FOXP3*, *IL2RA*), NMF2 (Th17-F; *RORC*, *CCR6*), NMF3 (Naive-F; *CCR7*, *BACH2*), NMF4 (Activation-F or Act-F; *DACT1*, *CDK6*), NMF5 (TregEff/Th2-F; *HLA-DR*s, *CCR10*), NMF6 (Tfh-F; *MAF*, *CXCR5*), NMF7 (Interferon-F or IFN-F; *OAS1*, *MX1*), NMF8 (Central Memory-F; *CRIP2*, *PLP2*), NMF9 (Thymic Emigrant-F; *SOX4*, *PECAM1*), NMF10 (Tissue-F; *JUNB*, *NFKBIA*) NMF11 (Th1-F; *GZMK*, *EOMES*)

At the major cluster L1 level, we did not detect any significant cell frequency changes in both CSF and PBMC (Supplementary Fig 7b and 8b). However, at the minor cluster L2 level, there was a significant reduction of CD4^+^ T effector memory (Tem) expressing T peripheral helper (Tph) markers (Tem -Tph; *PDCD1*^lo^*CXCR5*^+^) in both CSF (padj= 1.88 x 10^-2^) and PBMC (padj= 9.41x10^-6^) tissues after B cell depletion treatment (Fig 6c and d, Supplementary Fig 7c and 8c). In CSF alone, the frequency of CD4^+^ T central memory (Tcm) expressing T follicular helper (Tfh) markers (Tcm -Tfh; *PDCD1*^+^*CXCR5*^+^) was significantly reduced (padj=9.36x10^-3^) while the frequency of Tcm-Th0 was increased (padj=0.0472), suggesting a shift toward a naïve phenotype post- B cell depletion in the CNS (Fig 6c and d, Supplementary Fig 7c and 8c). We also observed a significant increase of CD4^+^ naive T cells expressing *SOX4* (Tnaive *SOX4*; *SOX4^+^PECAM1^+^*) (padj=4.29x10^-3^) in the blood, which is a recent thymic emigrant population^32^, indicating that the peripheral blood CD4^+^ T cell pool has been replenished by newly generated CD4^+^ T cells after the treatment (Fig 6d, Supplementary Fig 8c).

We then assessed the changes in gene program activity quantified by NMFproj in each L2 subpopulation. We observed a significant reduction in cell types for NMF6 (Tfh-Feature or Tfh-F; *MAF*, *CXCR5*) and NMF11 (Th1-F; *GZMK*, *EOMES*) post-treatment in both blood and CSF. (Fig 6e and f, Supplementary Data 1 and 2). Because NMF6 (Tfh-F) is predominantly high in Tcm-Tfh, Tem-Tph, and intermediate Treg (Treg Int), the decrease of NMF6 (Tfh-F) in both tissues indicates that B cell depletion treatment reduces their frequencies and potentially represses the repopulation of these CD4 subtypes quantitatively and qualitatively. An increase of NMF2 (Th17- F; *RORC*, *CCR6*) in the Tem population was observed in CSF, whereas NMF2 signature decreased in the blood (Fig 6e and f, Supplementary Data 1 and 2). We also observed increased NMF10 (Tissue-F; *JUNB*, *NFKBIA*) in blood after treatment (Fig. 6f, Supplementary Data 2). These observations suggested a redistribution of the CD4^+^ T subsets between the periphery and CNS. Altogether, these findings demonstrate that B cell depletion notably alters the CD4^+^ T cell compartment by reducing specific T cell populations such as Treg Int, Tcm-Tfh, and Tem-Tph and modifying effector gene expression profiles such as repression of NMF6 (Tfh-F) and NMF11 (Th1- F), which may be associated with its therapeutic efficacy in MS.

### B cell depletion increases suppressive Tregs

Loss of Treg function has been repeatedly observed in patients with MS^8,33^, and we showed MS susceptibility variant modulate can modulate Treg function^34^. We sub-clustered Tregs from scRNAseq PBMC data and compared L2 subpopulation frequencies to examine Treg alterations in more detail. We observed a significant decrease in naïve Tregs (Treg Naive) (*FOXP3, CCR7*, padj= 3.13x10^-2^) and Treg Int (*FOXP3*, *FCRL3*, padj=6.79x10^-3^) and an increase in effector Treg (Treg Eff) (*HLA-DR*s, *CD74*, padj=2.33x10^-3^) in post-treatment samples. (Fig 7a and b). We also examined gene expression differences in the whole Treg population and found that *HLA-DR*s and *CD74*, which are the markers of Treg Eff, were increased. In contrast, *FCRL3*, a marker of Treg Int, was decreased after treatment (Fig 7c, Supplementary Fig 9a and b). These data suggest that B cell depletion skewed the Treg function toward the effector phenotype. Next, we examined the potential mediators of myeloid-Treg interactions using ligand-receptor prediction analysis (Fig 7d). Since TNF is known to enhance the function of Tregs through interaction with TNFR2, which is abundant in Tregs^35,36^, we hypothesized that monocytes upregulated *TNF* expression following B cell depletion and promoted Treg expansion through TNFR2 signaling. We measured Treg frequency using flow cytometry in MS patient PBMC pre- and post- ocrelizumab treatment. We observed a significant increase in the frequency (p < 0.0001) of Tregs (Fig 7e) in post-treatment PBMC samples. TIGIT protein is highly expressed in Treg Eff^37^ (Supplementary Fig 9b and c) and has been shown to associate with increases in functional activity in human and mice^38,39^. Therefore, we measured the frequency of TIGIT expressing Tregs after B cell depletion and observed a significant increase post- ocrelizumab treatment (Fig 7f). In summary, these data indicate that B cell depletion corrects for the loss of immune regulation in MS by increasing Treg frequency and effector function.

**Fig 7:**
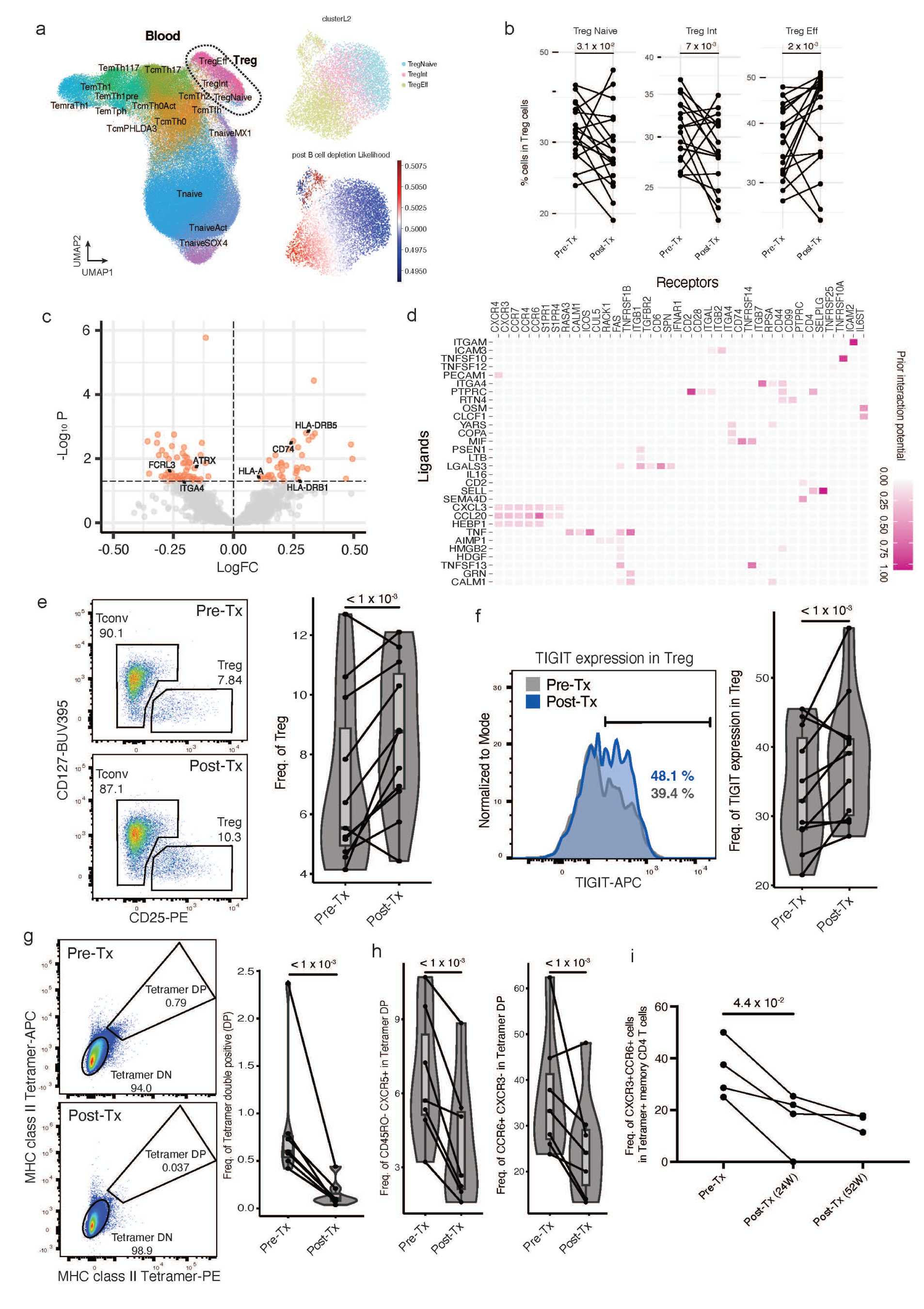
B cell depletion induces an increase in TIGIT^+^ Tregs and reduces autoreactive T cells. **a,** Visualization of Treg population extraction and changes after B cell depletion treatment. Predicted CD4^+^ T clusters and Tregs (dotted line) on UMAP plot (left). Re-embedding of extracted Tregs using UMAP (top right). B cell depletion treatment associated relative likelihood in Treg populations calculated using MELD (bottom right). **b,** Frequency changes of each subpopulation within the Treg group. **c,** Volcano plot depicting differentially expressed genes in Tregs, particularly highlighting genes encoding surface proteins. **d,** Heatmap displaying predicted interactions between myeloid cell-derived ligands (limited to genes differentially regulated with B cell depletion treatment) and Treg-derived receptors, weighted by prior interaction potential. **e,f,** Flow cytometry data of Tregs frequency **(e)** and TIGIT protein expression of Tregs **(f)** in MS patient PBMC (n=11) after B cell depletion treatment, **g,** Flow cytometry analysis of myelin tetramer- reactive CD4^+^ T cell frequency at pre-treatment and 6-month post-treatment timepoints (n=7). **h,** Tfh (CD45RA^-^CXCR5^+^) cells in tetramer-reactive CD4^+^ T cells and Th17 (CCR6^+^CXCR3^-^) cells in tetramer-reactive CD4^+^ T cells frequencies at pre-treatment and 6-month post-treatment timepoints (n=7). **i.** Longitudinal kinetic analysis of the frequency of autoreactive CCR6^+^CXCR3^+^ CD4^+^ T cells using flow cytometry at pre-treatment (n=4), 24-week post-treatment (n=4) and 52 week post-treatment (n=3) timepoints.

### B cell depletion decreases myelin tetramer binding CD4^+^ T cells

We and others have shown increases in the frequency of inflammatory myelin reactive T cells recognizing a number of myelin antigens, presumably as a consequence of epitope spreading, in the circulation of MS patients. We have demonstrated the utility of using a panel of MHC class II tetramers loaded with myelin epitopes from myelin basic protein (MBP), proteolipid protein (PLP), and myelin oligodendrocyte glycoprotein (MOG) to directly measure the frequency of autoreactive T cells in the disease^4,5,40,41^. To increase the accuracy of detecting autoreactive T cells, we used both PE- and APC-conjugated tetramers with the same myelin composition (Supplementary Fig 6b). We initially examined a cohort of patients pre- and post-B cell depletion and observed a marked decrease in the frequency of myelin PE- and APC-tetramer double positive (Tetramer DP) CD4^+^ T cells (Fig 7g). In these data, decreased frequencies of CD45RA- CXCR5^+^ cells and CCR6^+^CXCR3^-^ cells were observed in myelin tetramer binding CD4^+^ T cells (Fig 7h). Longitudinal kinetic analysis using fresh PBMC derived from a separate MS cohort (Supplementary Table 1) revealed significant decreases in the frequency of these autoreactive CCR6^+^CXCR3^+^ CD4^+^ T cells at 6 months post-treatment with no significant changes in CCR6^+^CXCR3^+^ cells in non-tetramer reactive CD4^+^ T cells. This trend continued through 52 weeks (Fig 7i). These results indicate that B cell depletion directly or indirectly regulates the frequency of presumably pathogenic myelin reactive T cells.

## Discussion

We performed an unbiased analysis of the immune landscape using scRNAseq after *in vivo* perturbation of the immune system with B cell depletion therapy in treatment-naïve MS patients. Ocrelizumab, a humanized anti-CD20 monoclonal antibody, induces a systematic removal of naïve and memory B cells with ∼98% efficacy in preventing new CNS lesions^3^. Here, we demonstrate that B cell depletion induces cell type-specific recovery of CSF macrophages and intermediate monocytes and reprograms them toward transcriptional phenotypes that resemble healthy donor macrophage and monocytes. Specifically, we identified a novel anti-inflammatory transcriptomic signature in post-treatment MS CSF macrophages, promoting the resolution of inflammation and maintain homeostasis with upregulation of CD83, TGFβ, and IL- 10. CD16^+^ monocytes showed the highest level of TNFα messenger RNA expression pre-treatment, and B cell depletion further increased TNFα protein expression in these monocytes. Furthermore, after B cell depletion, we observed shifts in T cell populations, including decreased frequencies of Tem-Tph and Treg Int, along with Th1- and Tfh-type gene programs. Notably, we found that B cell depletion increased TIGIT^+^ regulatory T cell frequency and decreased myelin tetramer-binding CD4^+^ T cell frequency in the blood compartment. Our study provides an extensive transcriptomic map of immunological changes through the simultaneous interrogation of the diverse immune populations perturbed by *in vivo* B cell depletion. This approach enables us to discover distinct cell-type specific mechanisms of action consistent with the remarkable efficacy of B cell depletion therapy in MS.

The changes in monocyte frequency after B cell depletion were unexpected and revealed the power of an unbiased systems approach in dissecting the multitudes of immune-regulatory mechanisms of a highly efficacious therapy. We hypothesize that the dichotomous roles of monocytes in the pathogenesis of MS are due to the nature in which myeloid phenotype and function are orchestrated by the metabolic requirements in surrounding tissue environments. Soluble factors and cytokines shape macrophage differentiation and activate molecular programs that either exacerbate or attenuate disease^42^. Systemic depletion of pathogenic B cells likely reduces the inflammatory states of multiple immune cell types in a feedback loop manner, and the subsequent recalibration of the cytokine milieu allows macrophages to differentiate in a steady-state environment and engage in disease-resolving transcription program to restore homeostasis. Given our study is limited to one post-treatment timepoint, it is possible that the changes we describe will normalize on the long term. Our ability to detect functional changes in the CSF compartment were limited by the small sample size, variable post-treatment time-points, and a small number of cells composing the CSF macrophages clusters compared to other immune populations. However, a previous study by Mrdjen et al. dissected changes in myeloid populations in the EAE model and observed CNS-specific alterations of macrophages and microglia phenotypes during the disease course^43^. Interestingly, they noted that border-associated macrophages (BAMs) share similar profiles with pDCs compared to other myeloid subsets. We similarly observed that increased abundance of CSF macrophages was accompanied by increased abundance of pDCs (Fig 1).

While the ontogeny of CSF macrophages has not been well elucidated, we posit that some of the enriched macrophage population could derive from blood monocytes as we observed some similarity in transcriptomic kinetics between enriched CSF macrophages and enriched CD16+ monocytes post- B cell depletion. With autoimmune exacerbations in patients with MS, peripheral blood monocytes receive inflammatory cytokine signals and cross the blood- brain barrier with T cells leading to CNS lesions^44,45^. While macrophages are the dominant cell type in these lesions, blood monocytes also extravasate into the CSF and differentiate into CSF macrophages^46^. However, it is critical to not hyperactivate CSF macrophages in the process of phagocytosing waste to prevent downstream immune activation of autoreactive lymphocytes^47^. With B cell depletion, the decrease in myeloid inflammatory cytokines enables CSF macrophages to receive pro-resolving signals to execute their homeostatic function in mediating tissue repair^48^. Our data suggest an anti-inflammatory phenotype in CSF macrophage after B cell depletion treatment and demonstrate that anti-CD20 depletion therapy restores macrophage gene signatures similar to those of healthy controls. Importantly, the inclusion of healthy donor CSF samples allows us to conclude that anti-CD20 treatment reprograms CSF macrophages toward a homeostatic/healthy cell state.

In the peripheral immune compartment, we uncovered a ubiquitous TNFα/NF_κ_B activation signature across a wide range of circulating immune cells post- B cell depletion. CD14^+^CD16^+^ monocytes have been shown to be a potent producer of TNFα^49^, and our study suggests that TNF is a pleiotropic cytokine that potentially exerts anti-inflammatory effects after B cell depletion. It has not escaped our attention that B cell depletion confers moderate clinical effectiveness in treating rheumatoid arthritis whereas anti-TNFα clearly worsen MS immunopathogenesis^50^. While the increased production of TNFα with a clinically effective treatment appears to be counter-intuitive, the beneficial role of TNFα in MS is substantiated by clinical trials results showing anti- TNFα treatment in patients led to significant worsening of disease activity^51^. Moreover, molecular dissection of MS risk allele rs1800693 located in the gene encoding the TNFR1 revealed the associated variant that codes for a soluble form of TNFR1, which mimics TNF blocking molecules^52,53^, again consistent with the observation that TNFα blockade leads to increased disease activity. In the context of chronic inflammation, this scenario where TNFα bears an anti-inflammatory role is reminiscent of the anti-tumor immune response. Specifically, in the tumor microenvironment, constant exposure to TNFα leads to immunosuppressive responses involving Tregs, B regulatory cells and myeloid-derived suppressor cells^54^, and blocking TNF leads to improved response to immune checkpoint blockade in an orthotopic melanoma mouse model^55^. Additionally, identifying the precise signaling events leading to the observed transcriptomic changes is challenging given that the main TNFα signaling pathway is through NF_κ_B, a highly ubiquitous signaling pathway. Thus, we cannot exclude other receptors signaling through NF_κ_B are participating to the transcriptomic alterations observed post-treatment. Nevertheless, our rich dataset provides a non-biased road map which can now be investigated in animal models to better elucidate the more detailed mechanisms associated with B cell depletion.

Several of the established cell annotation methods that we employed were unable to detect changes in the CD4 T cell compartment. The biologically relevant signals could potentially be obfuscated due to the small frequency of myelin-specific T cells. Applying reference mapping with finer granular reference and gene program quantification by NMF, we were able to detect B cell depletion- mediated modulation of various T cell subsets. While we were able to confirm the increase of effector Treg frequency both by transcriptomic and protein expression, it will be of interest to more precisely identify whether there are differences in specific myelin antigens and other antigens, as recent studies indicate EBV infection of B cells is associated with the onset of MS^56^.

Our T cell data suggests several mechanisms in which B cell depletion can lead to modulation of T cell functionality. One potential model is that TNFα from myeloid cells engages TNFR2 on Tregs leading to suppression of autoreactive T cells after B cell depletion in RRMS patients. Alternatively, as first shown by Lanzavecchia et al.^57^, B cells may be the key antigen presenting cell and their depletion may result in loss of autoreactive effector or memory T cells. Similarly, the decrease of MHC expression on myeloid cells may also support this hypothesis.

Rather than identifying one mechanism for the high clinical efficacy of B cell depletion in MS, our unbiased “*in vivo* perturbation” investigation uncovered a number of changes in the immune circuitry. This is perhaps not surprising as the genetic architecture of MS and other autoimmune diseases suggests that multiple pathways are involved in disease pathogenesis^58^. In summary, our unbiased systems analysis identified a series of immunoregulatory pathways induced by B cell depletion. We propose that for B cell depletion to be such a globally effective therapy in MS, essentially subverting the autoimmune attack of the CNS, several mechanisms of action must be involved to maintain homeostasis. It is likely that different immunosuppressive pathways become activated among patients, leading to marked decreases of autoreactive myelin reactive T cells in the blood compartment. Our analyses comparing immune cells in CSF and blood also highlights shared vs distinct changes across compartments, suggesting regulation of CNS homing mechanisms is affected by anti-CD20 therapies. Considering the potential of disease-mediating and homeostatic functions in the myeloid compartment, future analyses can be designed with a myeloid focus using fresh tissue for higher sensitivity in protein detection. Nevertheless, these datasets and observations provide a critical starting point that will require well-designed *in vitro* and *ex vivo* assays and appropriate animal models to fully elucidate the perturbational effects of B cell depletion on the functionality of the immune system.

## Methods

### Ethic Statement

This study was approved by the Institutional Review Board at Yale University. CSF and blood samples were obtained from healthy donors and MS patients with informed consent.

### Patient Cohorts

All patients had early onset relapsing remitting MS and had not been on previous immunomodulatory treatments. A small subset of patients had received IV solumedrol within 3 months of blood draw. Eighteen patients undergoing single cell RNA seq studies had CSF analysis prior to the initiation of treatment, and five of those subjects had repeat lumbar punctures, as outlined in the results section. A total of six age-matched healthy controls had lumbar punctures, and those results were previously reported^44^. An additional four subjects had flow cytometric analysis only. Patient characteristics are summarized in Supplementary Table 1.

### Sample preparation for scRNAseq

Fresh patient CSF samples were centrifuged, and cells were immediately processed using 10x Genomics 5Pv1 chemistry. Samples were collected prior to infusion of B cell depletion therapy. In the CSF sample cohort, four of the patients were administered ocrelizumab B cell depletion treatment, and one was treated with rituximab. Patient PBMCs were isolated from whole blood using Lymphoprep (STEMCELL) density gradient centrifugation. All patients were administered ocrelizumab B cell depletion in the PBMC cohort. Cryopreserved patient-matched pre-treatement and 6-month post-treatment PBMCs were thawed and processed within the same experimental batch using 10x Genomics 5Pv1 chemistry. For PBMCs, TCR libraries were generated along with the gene expression libraries.

### scRNAseq QC

PBMC and CSF libraries were sequenced at 20,000 read pairs per cell on Illumina NovaSeq instrument. Fastq files were processed using cellranger version 3.1.0 mapping to GRCh38 human reference genome. Alignment and quantitation were performed with the “cellranger count” command for each emulsion (using the 2020-A 10x genomics human reference), to generate unique molecular identifier (UMI) count matrices.

For CSF, Data QC was performed in R using the Seurat package. Low quality cells were filtered out based on mitochondria percent, UMI counts and number of features for individual samples. Samples were then merged, log10-transformed, and batch corrected using Harmony.

For PBMCs, we first filtered extreme outliers by excluding droplets with less than 1500 UMI counts, or less than 850 unique genes detected. As distribution of those parameters varied across emulsions, we median-centered the log10-transformed number of unique genes detected and removed low quality droplets with less than 1,100 unique genes detected or more than 2.5% mapping of UMI counts mapping to mitochondrial genes. We also removed potential doublets by filtering out droplets with more than 2,600 unique genes detected.

### scRNAseq analysis

#### Dimensionality reduction and clustering

For cells passing quality control, we normalized UMI counts by dividing each count by the total number of counts per cell. We then multiplied normalized counts by 10,000 and added a pseudo count of 1 before log-transformation. We computed the stabilized variance of each gene using the variance-stabilizing transformation (VST) and retained genes with stabilized variance > 1 for principal component analysis (PCA). Genes mapping to the T cell receptor (TCR), the B cell receptor (BCR) and the Y chromosome were excluded from PCA analysis. We computed the first 50 principal components (PCs) using a partial singular value decomposition method, based on the implicitly restarted Lanczos bidiagonalization algorithm (IRLBA), as implemented in the *Seurat* R package^59^. To correct for systematic differences across samples, we applied harmony integration^60^ to the first 50 PC loadings and retained 30 harmony-corrected PCs to build nearest neighbor graphs for visualization using Uniform Manifold Approximation and Projection (UMAP) (minimum distance = 0.5, spread = 10), and community detection using Louvain algorithm, as implemented in *Seurat*. We also computed a relative likelihood of cells being observed in specific experimental conditions using Manifold Enhancement of Latent Dimensions (MELD)^16^.

We embedded cells into 2 UMAP dimensions and applied Louvain algorithm. We annotated cluster cell types based on individual gene expression and the SingleR automatic annotation package using the MonacoImmuneData PBMC reference^61^. We tested for variation in clusters frequency in the CSF and in the blood separately by modelling the per-sample cluster frequencies using a beta distribution in a generalized linear model framework, as implemented in the *betareg* R package^62^.

#### Differential gene expression (DE)

For DE testing at the single cell level in CSF, we tested each gene individually using a generalized linear model approach as implemented in the *speedglm* R package, using a poisson distribution and a log link function^34^. We excluded low expression genes based on a UMI count per cell < 0.005, as well as ribosomal, mitochondrial, TCR, BCR. We evaluated differences in counts with treatment predictor, and a [number of UMIs detected] covariate to account for differences in library size. We then computed shrunk log fold changes using adaptative shrinkage methods implemented in the *ashr* R package^63^ (using a mixture of normal distributions) and FDR (Benjamini & Hochberg method) across all genes tested. Genes with FDR < 0.05 were considered differentially expressed. For differential expression testing at the single cell level in PBMCs, we used a negative binomial mixed linear model as implemented in the NEBULA package^28^. We then used shrunken log fold changes as a ranking metrics to run geneset enrichment analysis (GSEA).

#### Treg volcano and ligand-receptor analyses

Volcano plot displaying differential expression analysis performed using *nebula* comparing pre- and post-treatment Treg population. Among genes with differential expression (BH < 0.05) and average expression >0.1, those that encode surface proteins (based on Cell Surface Protein Atlas surfaceome protein database) were selectively labeled^64^. NicheNet (*nichenetr)*^65^ was used to identify predicted ligand-receptor interactions between myeloid populations and Tregs, will a particular focus on potential ligands that are differentially regulated in myeloid cells with B cell depletion treatment. Tregs were selected as the “receiver cell type”, including all expressed genes as potential receptors. Myeloid cells were selected as the “sender cell type”, limiting the set of potential ligands to the combined list of genes differentially expressed with B cell depletion treatment in myeloid cell clusters (see *NEBULA* analysis). Predicted ligand-receptor interactions were displayed as a heat map in which ligands (expressed by myeloid cells) were plotted against receptors (expressed by Tregs) and weighted by prior interaction potential.

#### CD4^+^ T cell automatic labeling and quantification of gene programs

The PBMC and CSF data were processed using the pipeline developed in the previous study^31^ to assign CD4**^+^** T cell clusters. This pipeline employs Azimuth^37^ for the extraction of CD4**^+^** T cells and uses Symphony^66^ for predicting CD4**^+^** T cell clusters. For interpretability, ’Treg Act’ has been renamed to ’Treg Int’ from the original literature. We tested for variation in cluster frequency by modeling the per-sample cluster frequencies using a beta distribution in a generalized linear model framework, as implemented in the *betareg* R package. For the assessment of TIGIT protein expression, we used CITE-seq data from PBMCs deposited in GSE164378 and performed reference mapping using the pipeline. Additionally, a 12-dimensional qualitative evaluation was conducted on the extracted CD4**^+^** T cells using NMFproj^67^. We applied a generalized linear model to assess feature changes per cluster^31^.

### Flow cytometry analysis

Frozen PBMCs were used for flow cytometry validation, except longitudinal myelin tetramer staining were performed on fresh PBMCs (n=4). Patient peripheral blood mononuclear cells were stained with a ViaKrome 808 Fixable Viability dye following the manufacturer’s instructions. Cells were then labeled with surface antibodies for 30 min at 4°C. For intracellular staining, cells were fixed and permeabilized with BD Cytofix/Cytoperm Buffer (BD Biosciences) for 10 min at room temperature, then washed with phosphate-buffered saline. Intracellular proteins were stained in permeabilization buffer (eBioscience) for 30 min at 4 °C. Antibody details are provided in Supplementary Table 2. For TNF staining, monocytes were enriched from cryopreserved PBMCs using EasySep™ Human Monocyte Enrichment Kit without CD16 Depletion Kit (STEMCELL technologies). Enriched monocytes were stimulated with 100ng/ml LPS for 4h at 37 °C before staining. To investigate myelin tetramer reactive T cells, APC- or PE- conjugated tetramers which were composed by DRB1*15:01 (loaded with MBP, MOG and PLP) or DRB1*04:01 (loaded with MOG and PLP) were used^4,41^. Myelin tetramers were incubated with cells for 30 min at 37°C before staining with antibodies. Cells were acquired on a BD Symphony flow cytometer with FACSDiva (BD Pharmingen) and data were analyzed with FlowJo software v.10 (Treestar).

### Peptide loading

Biotinylated monomers were diluted to a concentration of 0.5 mg/mL with 0.1 M phosphate buffer and incubated with 0.4 mg/ml of at 37°C for 72 h in the presence of 2.5 mg/ml n-Octyl β-D-glucopyranoside (OG) and 1 mM Pefabloc SC (Sigma–Aldrich, St. Louis, MO). Peptide loaded monomers were subsequently conjugated into tetramers using R-PE streptavidin (ThermoFisher Scientific, Waltham, MA) or fluorochromes of interest at a molar ratio of 8:1. Myelin peptide sequences are listed in the Supplementary Table 3.

### Data and code availability statement

All raw scRNAseq data generated in this study will be deposited on dbGAP upon acceptance. All code used for genomics analysis will be available on github and figshare, along with intermediate analysis files upon acceptance.

## Acknowledgement

The authors thank Rahul Dhodapkar for helpful discussions on computational data analysis. Chuan He for technical support and Kathryn Miller-Jensen for experimental input and discussion. This work was supported by NIH grants (P01 AI073748, U19 AI089992 U24 AI11867, R01 AI22220, UM 1HG009390, P01 AI039671, P50 CA121974, R01 CA227473) and Race to Erase MS to D.A.H. and National MS Society to D.A.H. and M.R.L. and grants from Genentech and F. Hoffmann-La Roche Ltd as part of Integrative Neuroscience Collaborations Network.

