## supplementary figure for "Systems Analysis of Immune Changes after B-cell Depletion in Autoimmune Multiple Sclerosis"

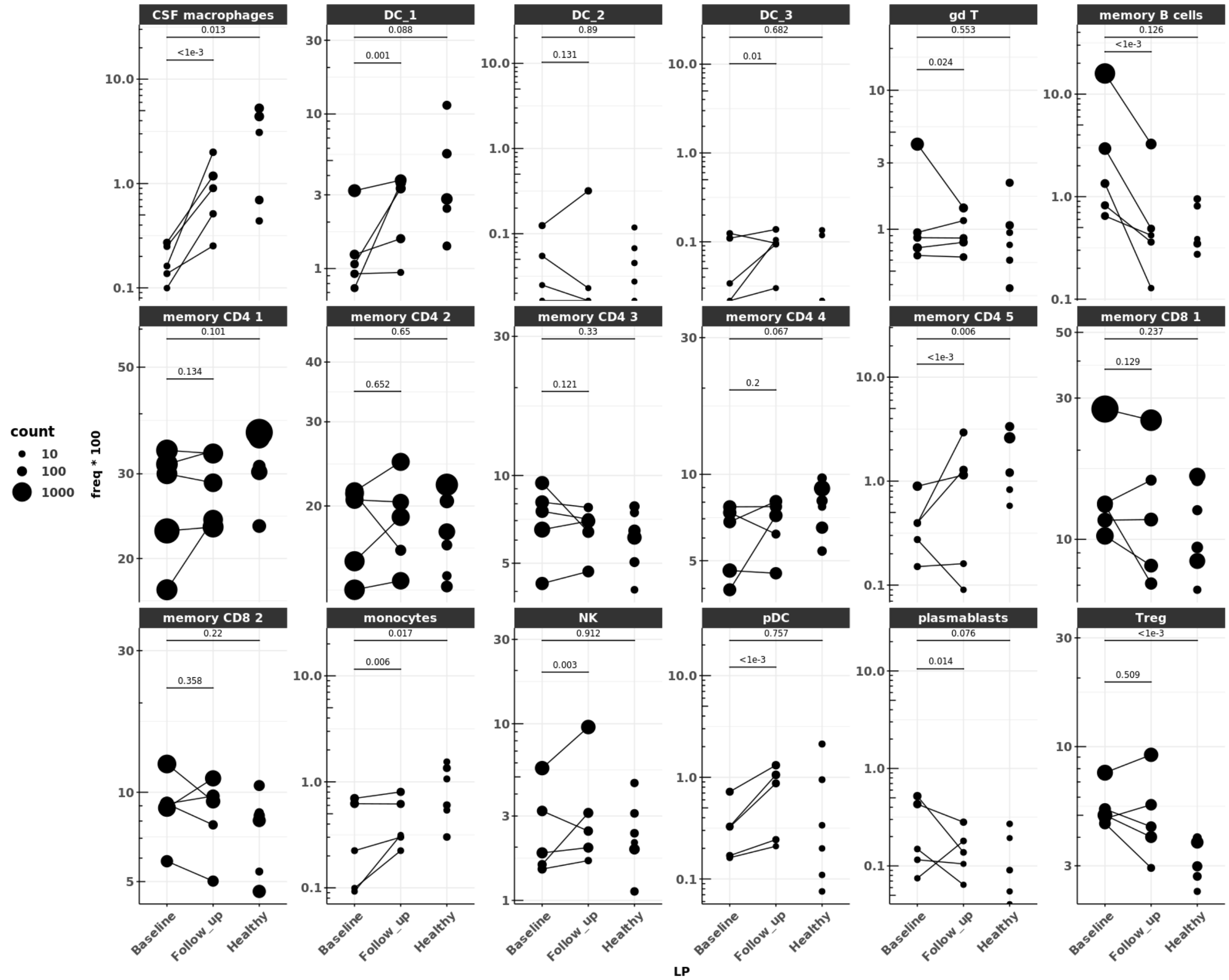


**Supplementary figure 1: Cluster frequency changes in the CSF.**

Per sample cluster frequencies in baseline MS patients with paired follow up samples (n=5) and healthy donors (n=6)

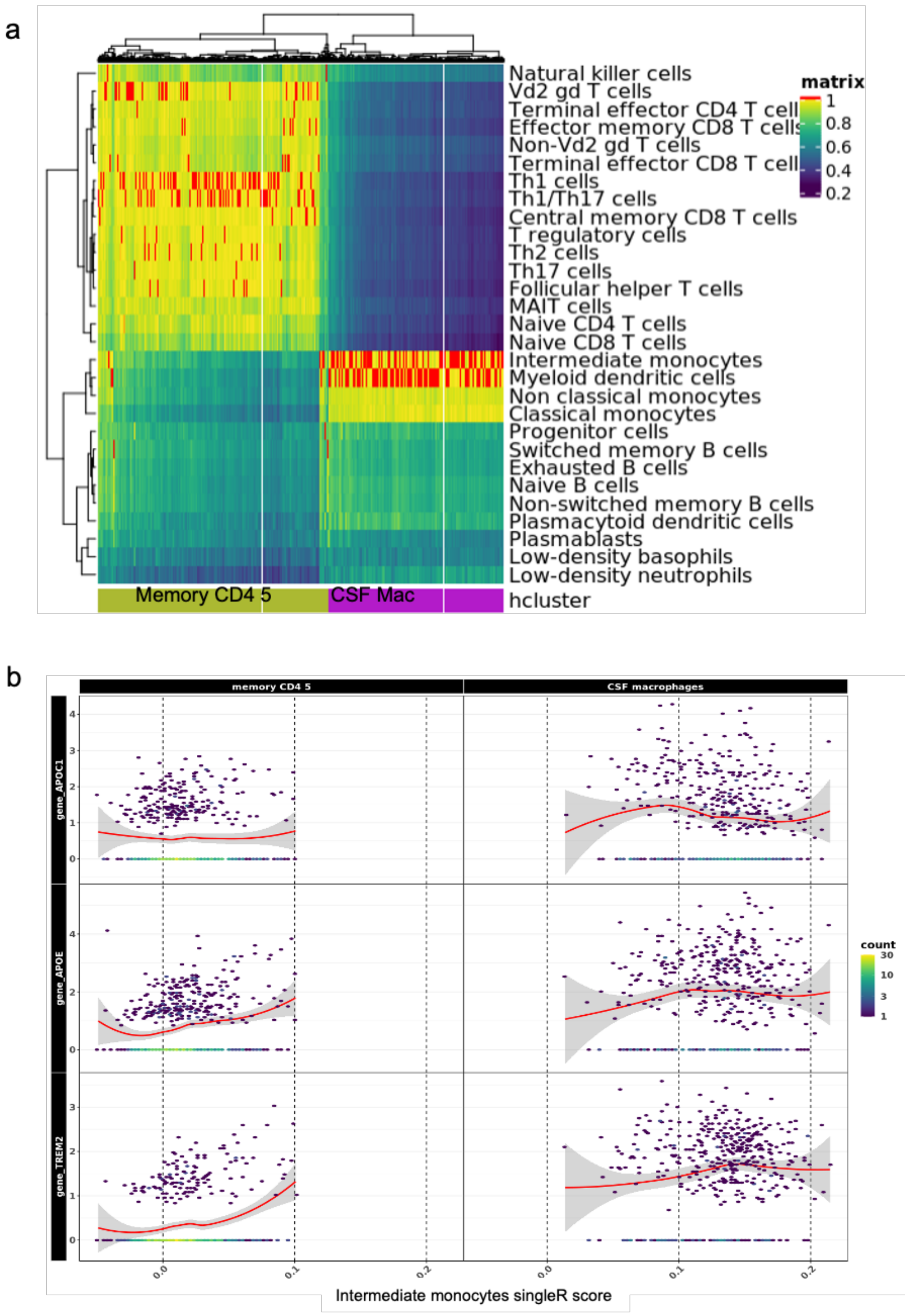


**Supplementary figure 2: Common CSF signature between macrophage and T cell clusters.** **a,** Community based clustering (Louvain) on shared nearest neighbor graph led to a cluster with mixed lineages (T cells and myeloid cells), based on singleR scoring against the Monaco reference. Separation of the two lineages by hierarchical clustering let us define two clusters for downstream analysis. **b,** 'Intermediate monocytes' singleR score separates each lineage, but both lineages express similar gene signature associated with CSF residency (*APOC1, APOE, TREM2*).

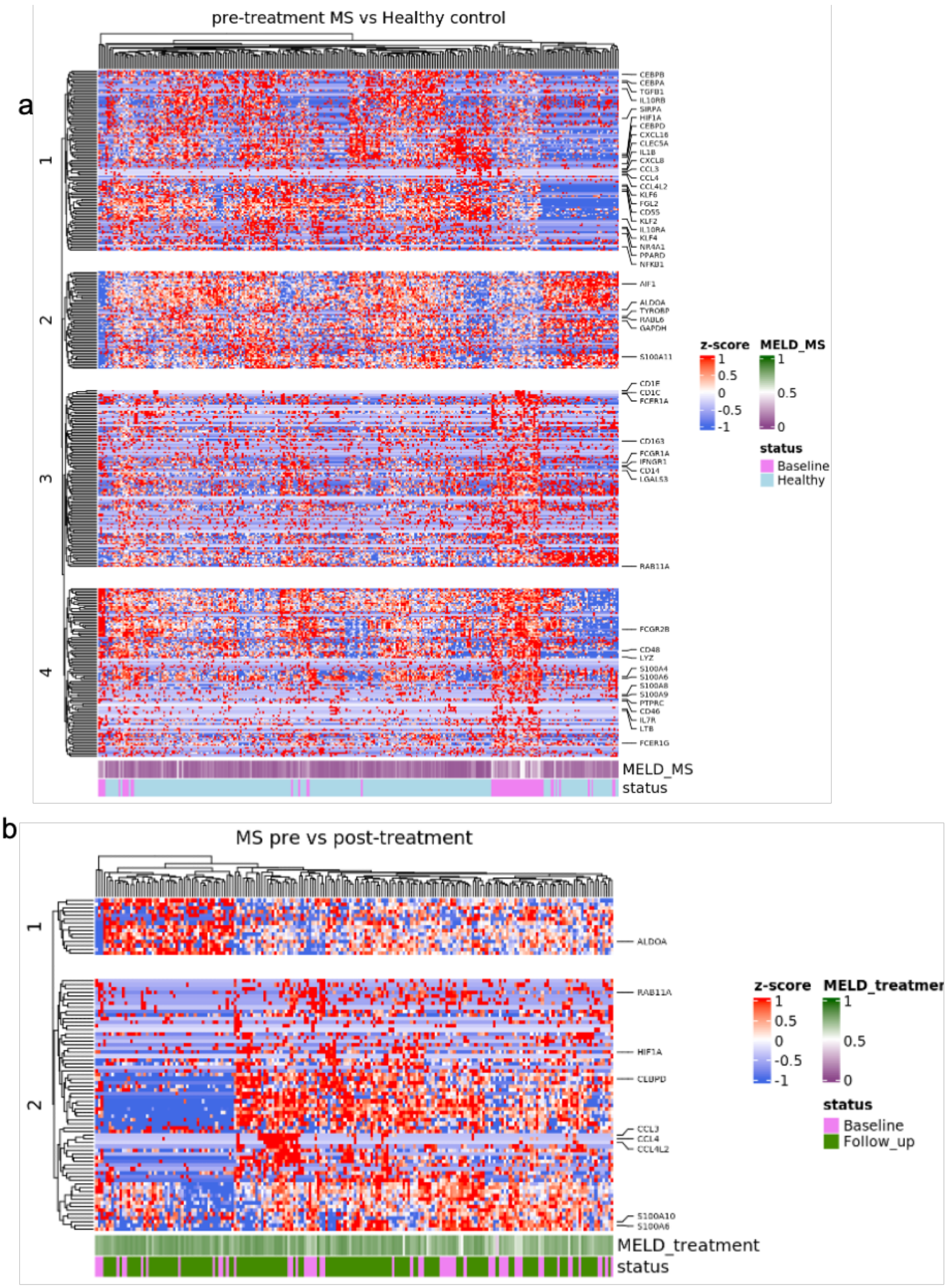


**Supplementary figure 3: Transcriptomic changes in CSF macrophapes.**
Heatmap of differentially expressed genes in the 'Mac 1' CSF macrophage cluster for MS baseline vs healthy donor **(a)** and MS follow up vs baseline **(b)**.

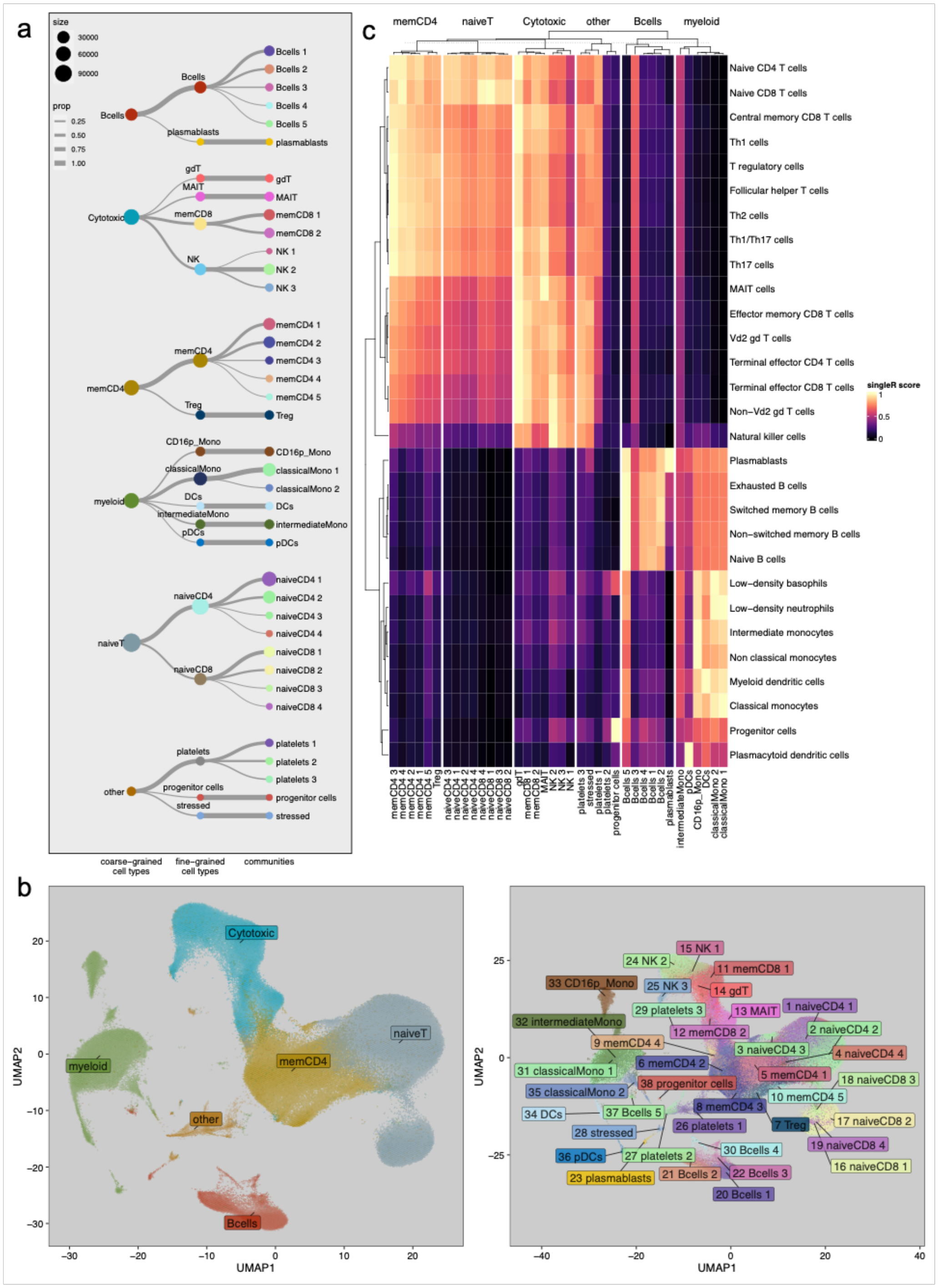

**Supplementary figure 4: Community detection and aggregation of PBMC single cell RNAseq dataset.**

**a,** initial community detection resolution (coarse-grained cell type) and subsequent nested community detection (community). Communities were then merged into fine-grained cell types. **b,** UMAP embedding annotated with coarse-grained cell types assignments and communities assignments. **c,** singleR scores heatmap for fined-grained cell types, grouped by coarse-grained cell types.

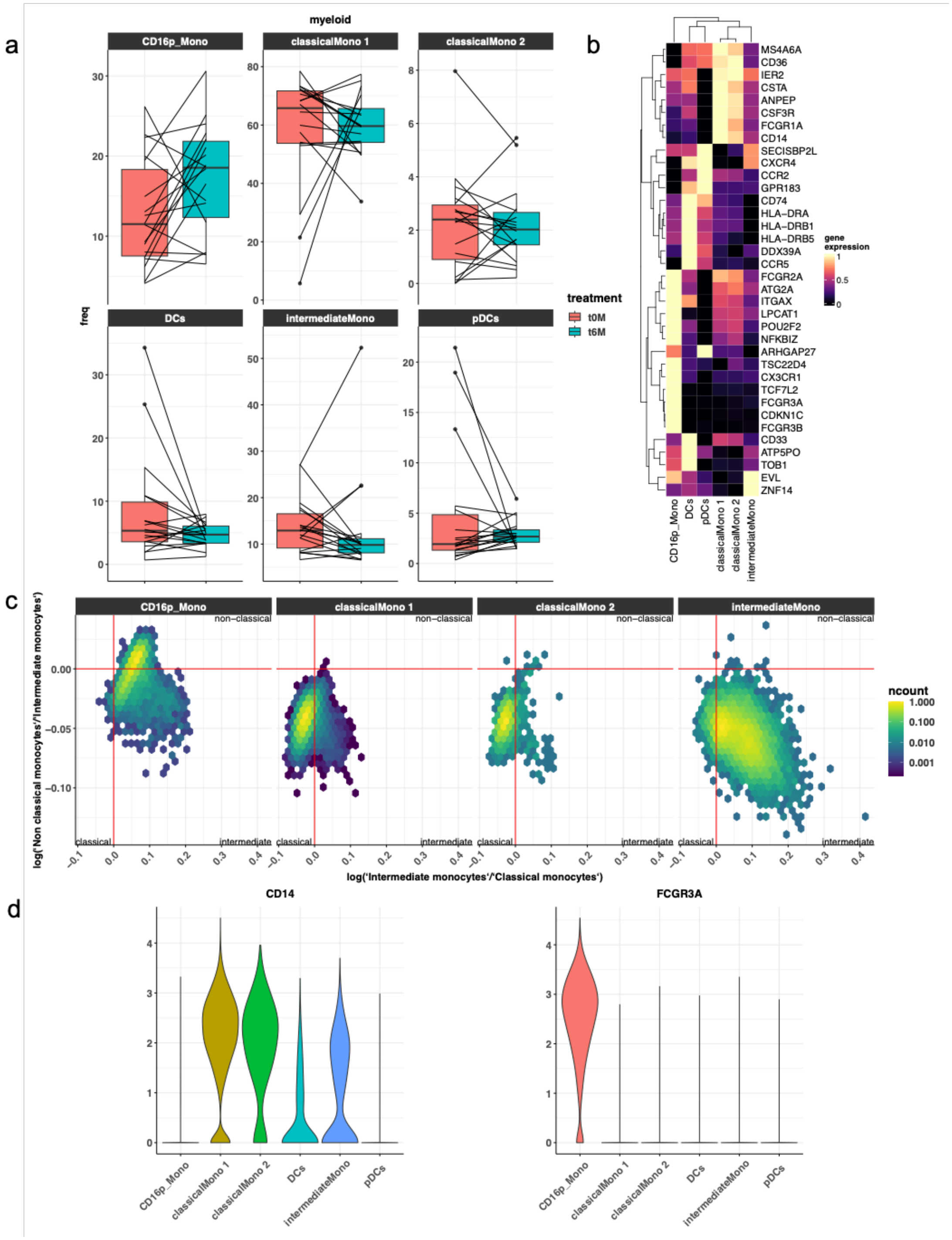

**Supplementary figure 5: Characterisation of myeloid clusters in PBMC dataset.**
**a,** frequency changes in myeloid clusters in MS follow up vs baseline samples. **b,** heatmap of myeloid cell type gene markers across fine-grained cell types. **c,** singleR monocytes subtypes score comparison across clusters defines 3 quadrants to differentiate between classical, intermediate, and non-classical subsets. CD16+ monocytes cluster presents with a mixture of intermediate and non-classical cells. **d,** RNA expression of CD14 and FCGR3A (encoding CD16) in myeloid clusters.

a

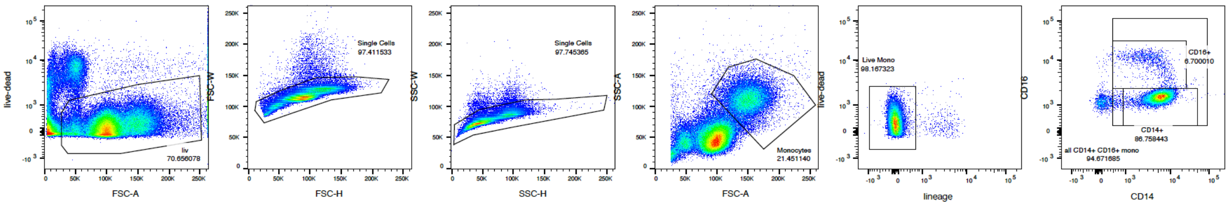


b

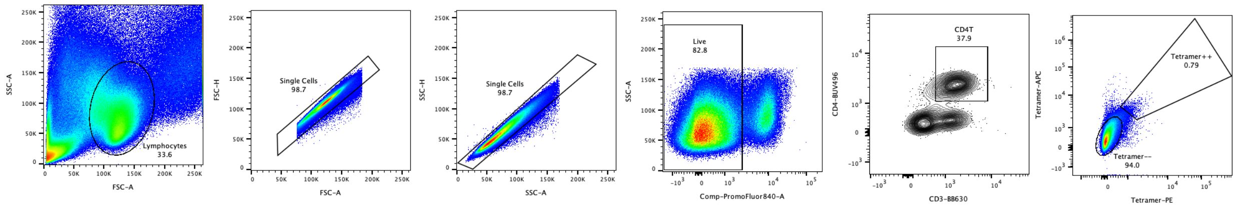


**Supplementary figure 6: Flow cytometry gating strategy.**

**a,b,** Flow cytometry images showed the gating strategy for monocytes **(a)** and lymphocytes **(b)**.

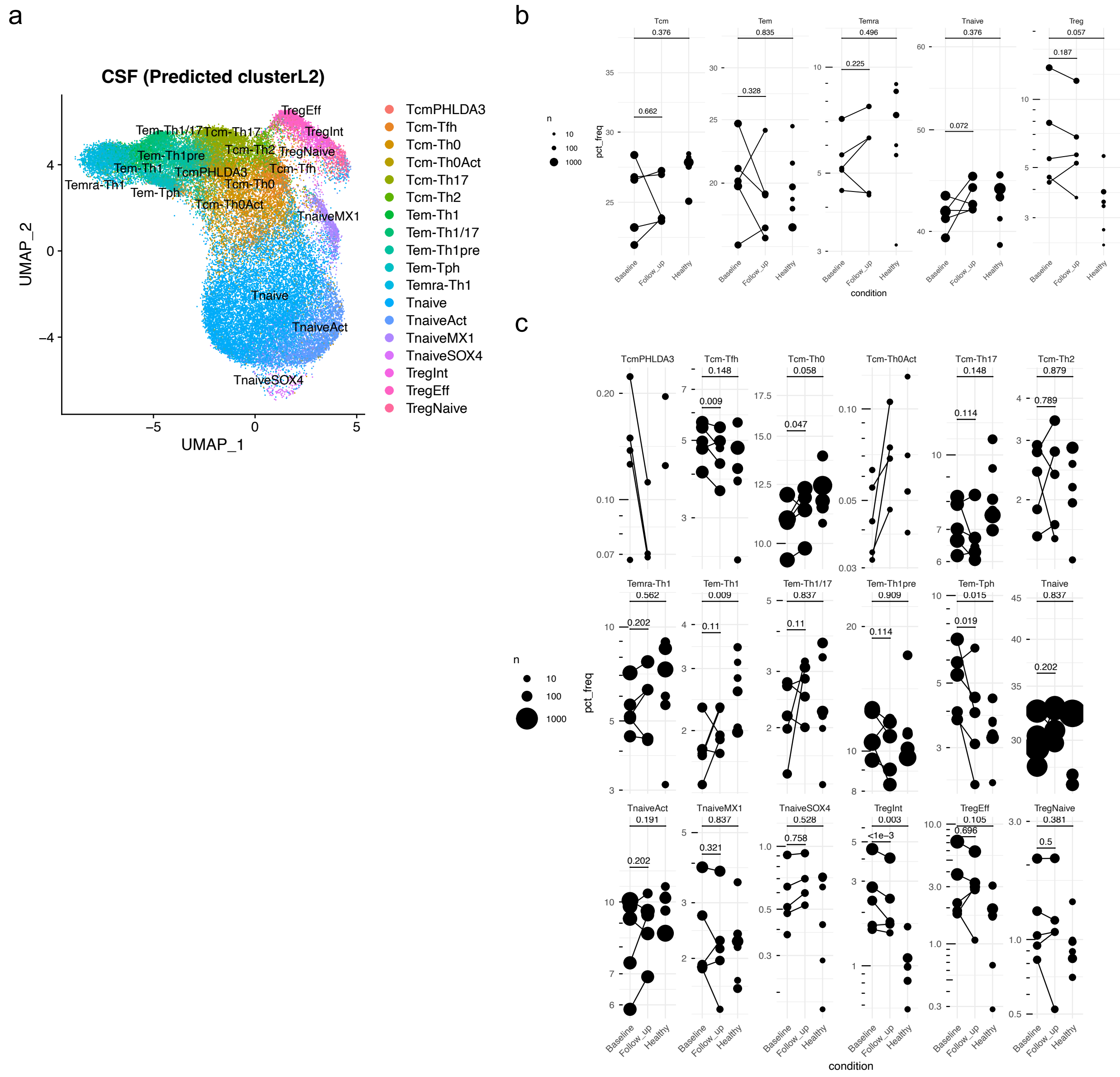


**Supplementary Figure 7: CD4+ T cell alterations in CSF**

**a,** Inferred CSF CD4$^{+}$ T cell clusters on UMAP plot. The clusters were assigned to a detailed cluster (cluster L2) level. **b, c,** CSF CD4$^{+}$ T cell frequency changes after anti-CD20 treatment in cluster L1 level **(b)** and cluster L2 level **(c)**. Coefficients of cell frequency change per cluster L2 quantified GLM (method) are visualized on the UMAP plot (left). CD4$^{+}$ T cluster frequency pre- and post-B cell depletion therapy (right).

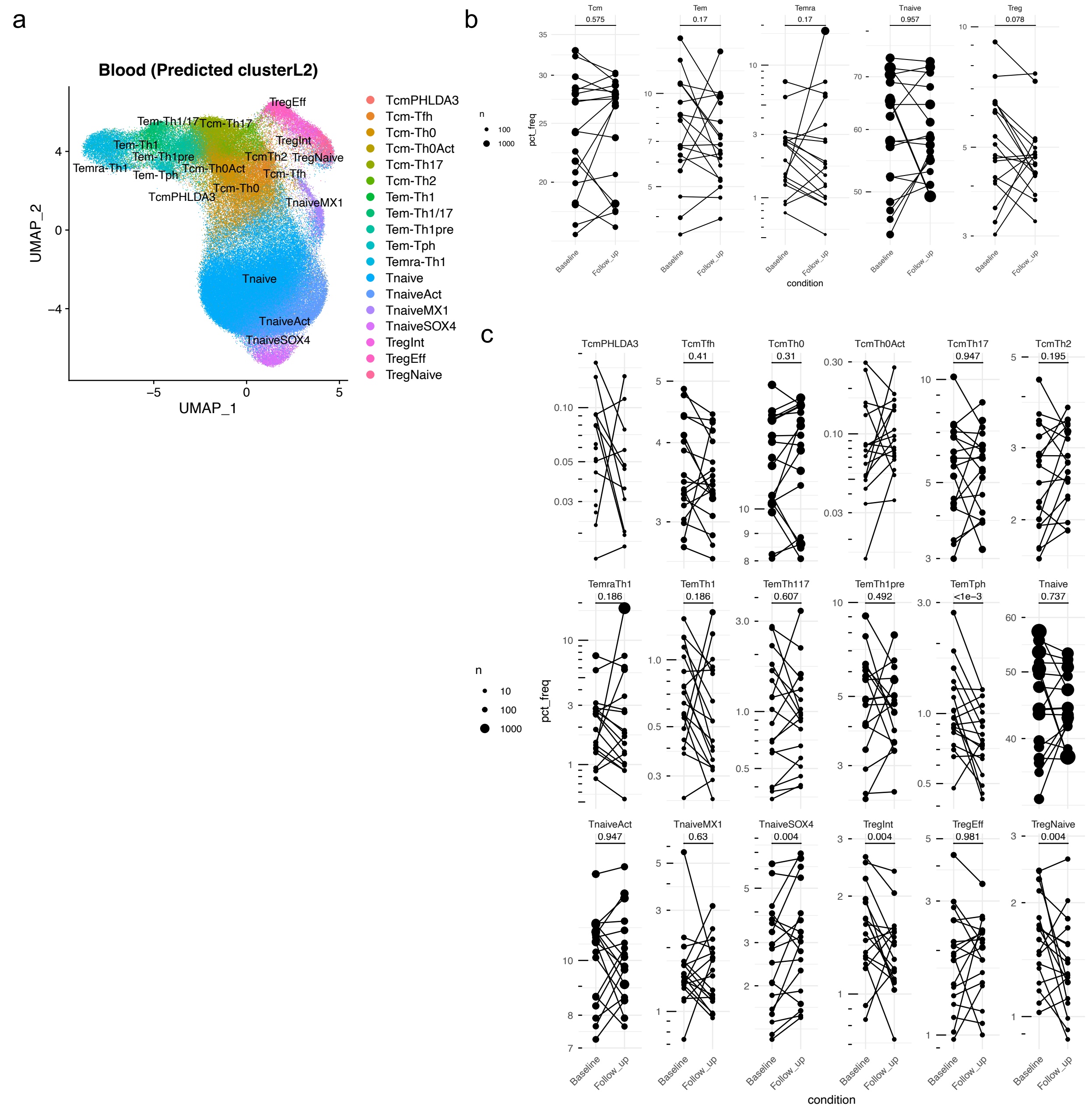


**Supplementary Figure 8: CD4+ T cell alterations in blood**

**a,** Inferred blood CD4$^+$ T cell clusters on UMAP plot. The clusters were assigned to a detailed cluster (cluster L2) level. **b, c,** Blood CD4$^+$ T cell frequency changes after anti-CD20 treatment in cluster L1 level **(b)** and cluster L2 level **(c)**. Coefficients of cell frequency change per cluster L2 quantified GLM (method) are visualized on the UMAP plot (left). CD4$^+$ T cluster frequency pre- and post-B cell depletion therapy (right).

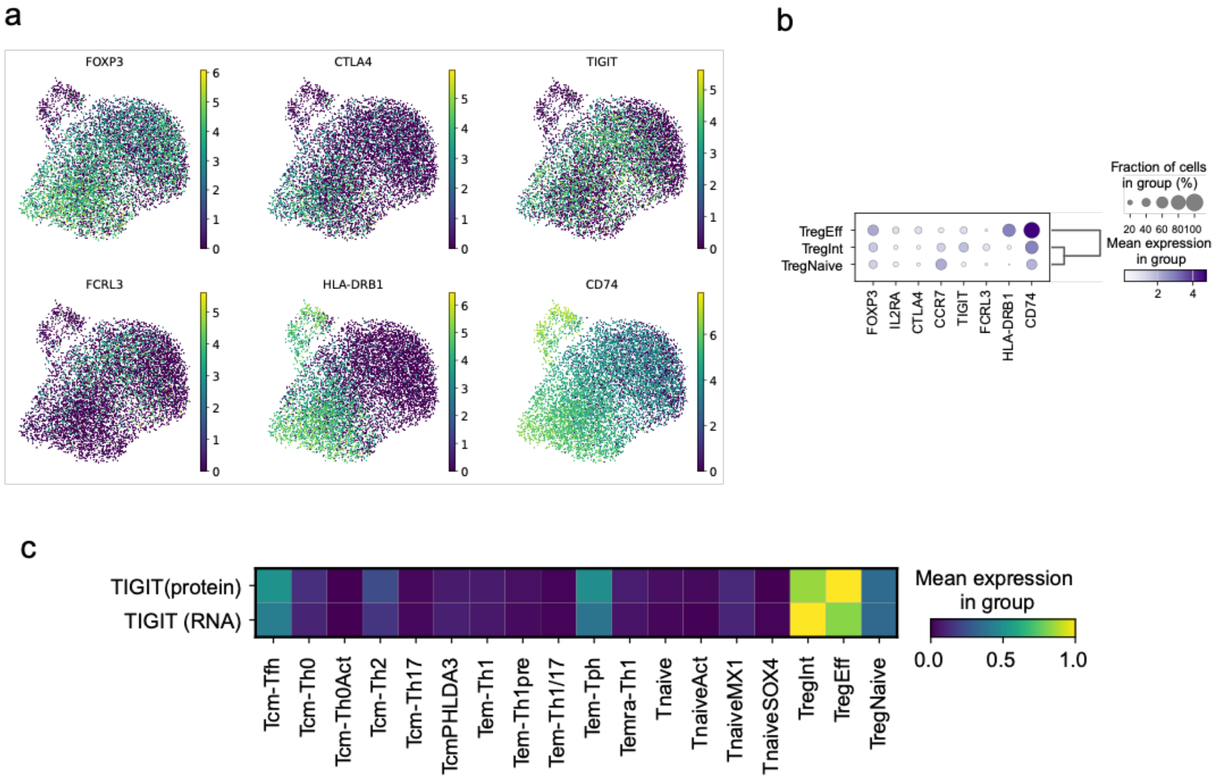


**Supplementary Figure 9: Detailed Treg characteristics**

**a,b,** Treg marker gene expression. Blood Treg cells were extracted and shown using UMAP **(a)** or dot plot **(b)**. **c,** Heatmap showing protein and RNA expression of TIGIT in cluster L2 clusters. CITE-seq data (GSE164378) was used for the visualization.
